# Transposon-encoded nucleases use guide RNAs to selfishly bias their inheritance

**DOI:** 10.1101/2023.03.14.532601

**Authors:** Chance Meers, Hoang Le, Sanjana R. Pesari, Florian T. Hoffmann, Matt W.G. Walker, Jeanine Gezelle, Samuel H. Sternberg

## Abstract

Insertion sequences (IS) are compact and pervasive transposable elements found in bacteria, which encode only the genes necessary for their mobilization and maintenance. IS*200*/IS*605* elements undergo ‘peel-and-paste’ transposition catalyzed by a TnpA transposase, but intriguingly, they also encode diverse, TnpB- and IscB-family proteins that are evolutionarily related to the CRISPR-associated effectors Cas12 and Cas9, respectively. Recent studies demonstrated that TnpB-family enzymes function as RNA-guided DNA endonucleases, but the broader biological role of this activity has remained enigmatic. Here we show that TnpB/IscB are essential to prevent permanent transposon loss as a consequence of the TnpA transposition mechanism. We selected a family of related IS elements from *Geobacillus stearothermophilus* that encode diverse TnpB/IscB orthologs, and showed that a single TnpA transposase was active for transposon excision. The donor joints formed upon religation of IS-flanking sequences were efficiently targeted for cleavage by RNA-guided TnpB/IscB nucleases, and co-expression of TnpB together with TnpA led to significantly greater transposon retention, relative to conditions in which TnpA was expressed alone. Remarkably, TnpA and TnpB/IscB recognize the same AT-rich transposon-adjacent motif (TAM) during transposon excision and RNA-guided DNA cleavage, respectively, revealing a striking convergence in the evolution of DNA sequence specificity between collaborating transposase and nuclease proteins. Collectively, our study reveals that RNA-guided DNA cleavage is a primal biochemical activity that arose to bias the selfish inheritance and spread of transposable elements, which was later co-opted during the evolution of CRISPR-Cas adaptive immunity for antiviral defense.

## INTRODUCTION

Some of the largest and most rapid forms of genetic diversification result from the activity of transposable elements (TEs), which are present in all domains of life and function as major drivers of genome evolution^1,2^. The selfish spread of diverse TEs within and between genomes relies on an eclectic mix of nucleic acid-metabolizing enzymes that act on both the RNA and DNA level, with classical retrotransposons (Class I) and DNA transposons (Class II) being mobilized by reverse transcriptases and transposases, respectively^3^. TE propagation often poses a severe fitness cost on host cells, leading to the emergence of regulatory pathways that manage this genetic conflict, but TEs and host cells also engage in more complex interactions that can result in mutualistic cooperation or even active cooption of TE sequences and genes^4,5^. Indeed, many essential cellular processes evolved directly from mobile genetic elements (MGEs) — including mRNA intronic splicing^6,7^, telomere maintenance^8^, immunoglobulin diversification^9^, and spacer acquisition^10^ — highlighting the molecular opportunities afforded by natural exaptation and domestication of transposon-encoded genes^4,11^. Recent research efforts similarly showcase the technological utility of these genes for DNA cleavage/joining reactions and genome engineering applications^12^.

Bacterial and archaeal adaptive immune systems encoded by CRISPR-Cas loci represent a particularly compelling example where recurring gene exchange between host and transposon has played a pervasive evolutionary role^11^. CRISPR arrays are themselves the products of sequential DNA integration reactions catalyzed by an enzyme called Cas1, which is a homolog of transposases found within so-called Casposon TEs, suggesting that the early origins of adaptive immunity required the cooption of *cas1* genes by proto-CRISPR-Cas systems^13^. In the more recent evolutionary past, mature CRISPR-Cas systems have been stolen and repurposed by diverse MGE types, including plasmids^14^, phages^15^, and transposons^16-18^, where they are thought to promote selfish spread through the targeted degradation and exclusion of other MGE types, or via RNA-guided DNA transposition^19^. And last but not least, the hallmark RNA-guided DNA cleaving enzymes that define Type II and Type V CRISPR-Cas systems — Cas9 and Cas12 — are evolutionarily related to IscB and TnpB enzymes encoded by IS*200*/IS*605*-family TEs^20,21^. Thus, although CRISPR-Cas systems canonically offer their hosts immunity from foreign MGEs, their very genesis is intimately linked with gene domestication events that provided critical and novel biochemical capabilities^11^.

Bacterial TEs that encode only the gene(s) necessary for transposition are referred to as Insertion Sequences (ISs), and IS elements within the IS*200*/IS*605* superfamily are mobilized by an HUH endonuclease-superfamily transposase known as TnpA, which contains a single conserved catalytic tyrosine residue (Y1) required for transposition^22^. Whereas autonomous IS*200* elements encode only *tnpA*, related TEs within the IS*605* family that bear conserved left end (LE) and right end (RE) sequences sometimes encode *tnpA* alongside an accessory nuclease gene known as *tnpB* or *iscB*, and more often encode *tnpB/iscB* alone^23^. Early studies demonstrated that *tnpB* is dispensable for *tnpA*-mediated transposition, indicating that most IS*605*-family elements function as non-autonomous TEs, but they failed to convincingly reveal a biological function for TnpB^24,25^. More recently, the Zhang and Siksnys groups reported that TnpB and IscB nucleases both function as RNA-guided DNA nucleases that use noncoding RNAs encoded by the IS element itself^26,27^. These remarkable findings demonstrated that the ability for single-effector, CRISPR-associated proteins to target and unwind double-stranded DNA (dsDNA) using an RNA guide first arose in bacterial transposons. However, whether and how this molecular property plays a role within the transposition pathway of these elements has remained enigmatic (**Fig. 1a**).

**Figure 1.**
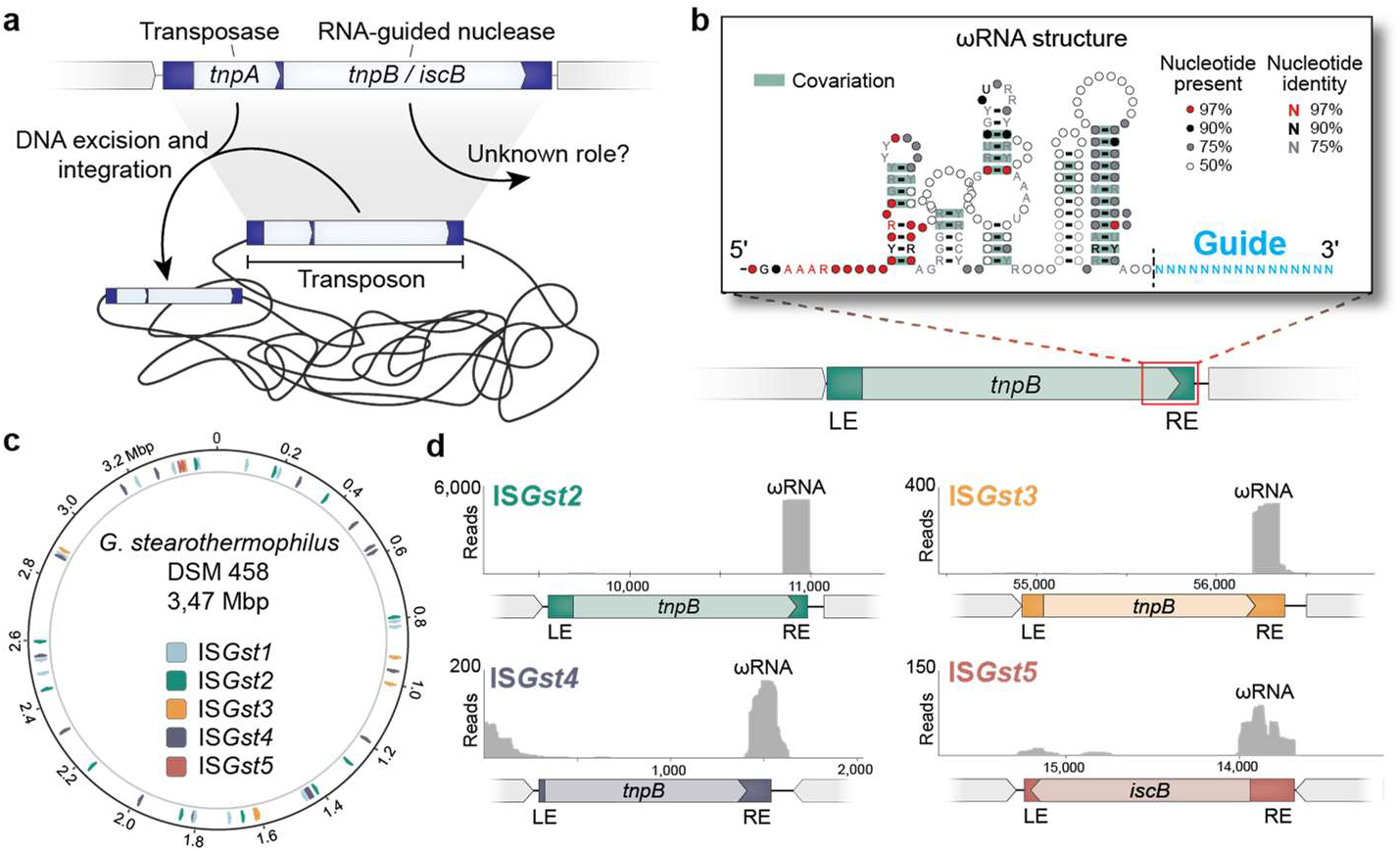
Pervasive distribution of IS*200*/IS*605*-like elements in *G. stearothermophilus*. **a,** Schematic of a representative IS*200/*IS*605* element. *TnpA* encodes a Y1-family tyrosine transposase responsible for DNA excision and integration; *tnpB/iscB* encode RNA-guide nucleases whose biological roles are unknown. **b,** Schematic of a non-autonomous IS element encoding TnpB and its associated overlapping ωRNA; a structural covariation model is shown in the inset. The green rectangle indicates the transposon boundaries, and the guide portion of the ωRNA is shown in blue. LE and RE, transposon left end and right end. **c,** Genome-wide distribution of IS*200*/IS*605*-like elements in *G. stearothermophilus* strain DSM458. Five distinct families are shown (IS*Gst1-5*), based on sequence similarity of transposon ends and nuclease encoded. **d,** Read coverage from small RNA-seq data of *Gst* strain ATCC 7953^36^, demonstrating expression of putative ωRNAs from each of the indicated IS*Gst* families. TnpB-associated ωRNAs are encoded within/downstream of the ORF, whereas IscB-associated ωRNAs are encoded upstream of the ORF.

Here we show that retention of IS*200*/IS*605* transposons at the donor site after DNA excision, and thus long-term transposon survival, relies critically on DNA cleavage by TnpB/IscB nucleases. By exploiting transposon-encoded guide RNAs to specifically recognize excision products and generate genomic DNA double-strand breaks, TnpB/IscB trigger host-mediated recombination that reinstalls the transposon, thus providing an alternative pathway to achieve proliferative transposition that we term ‘peel-and-paste/cut-and-copy.’ We furthermore uncovered a striking evolutionary convergence, in which both the transposase and nuclease enzymes encoded by IS*200*/IS*605*-family elements recognize overlapping transposon-adjacent motifs (TAMs) that abut the DNA integration site, ensuring a tight molecular coupling between the otherwise distinct processes of DNA excision, DNA insertion, and DNA cleavage. Beyond uncovering an elegant mechanism of transposon maintenance, this work advances our understanding of the biological function of a large family of RNA-guided nucleases that are encoded within diverse transposable elements found in all domains of life^28-30^.

## RESULTS

### *G. stearothermophilus* encodes diverse TnpB/IscB homologs

We set out to explore the evolutionary diversity of TnpB and IscB proteins — which, like Cas12 and Cas9, contain RuvC or RuvC and HNH nuclease domains, respectively — and identify conserved genes and sequence elements within their genetic neighborhood, as an entry point to investigating their function^20^. We first mined the NCBI NR database for TnpB/IscB homologs and built phylogenetic trees that highlight the diversity of both protein families (**Extended Data Fig. 1a,d**)^20,26^. When we extracted flanking genomic regions, we identified only a sporadic association with Y1 tyrosine transposases, with ∼25% of all *tnpB* genes containing an identifiable *tnpA* nearby, indicative of autonomous transposons. Interestingly, *iscB* genes were much less abundant than *tnpB* and rarely associated with *tnpA* (∼1.5%). This suggests that the vast majority of *tnpB/iscB* genes are encoded within transposons lacking *tnpA*, suggesting a non-autonomous function that would require transposases encoded elsewhere to mobilize them *in trans* (**Extended Data Fig. 1a,d**). *TnpB* but not *iscB* genes were also found associated with an unrelated serine resolvase (also denoted *tnpA*) that is a hallmark of IS*607*-family transposons, albeit at a much lower frequency (∼8%), consistent with previous observations.^31,32^ (**Extended Data Fig. 1d**).

In our initial analyses, we also noticed a conspicuous, highly conserved intergenic region upstream of *iscB* that was bounded by the transposon right end (RE), which bore striking similarity to a non-coding RNA termed HEARO previously described by Breaker and colleagues^33^. Similarly, non-coding RNAs (termed sotRNAs) were detected downstream of *tnpB* genes in Halobacteria by Koide and colleagues^34,35^, and elegant work from the Zhang and Siksnys groups recently revealed that both IscB and TnpB use these transposon-encoded RNAs — referred to hereafter as ωRNAs — as guides to direct cleavage of complementary dsDNA substrates, in a mechanism analogous to Cas9 and Cas12^26,27^. We generated covariation models for TnpB- and IscB-specific ωRNAs, which revealed the conserved secondary structural motifs characteristic of both guide RNAs (**Fig. 1b, Extended Data Fig. 1b**), and used these models to demonstrate the tight genetic linkage between *tnpB/iscB* genes and flanking ωRNA loci (**Extended Data Fig. 1a,d**). In order to investigate whether ωRNA production might be sensitive to local genetic context, we analyzed the orientation of genes upstream of *iscB* throughout the diverse members in our phylogenetic tree and observed a strong bias for genes encoded in the same orientation (**Extended Data Fig. 1c**). Since IscB-specific ωRNAs comprise a constant scaffold sequence derived from the transposon RE, joined by a 5’-adjacent guide region encoded outside of the transposon boundary, ωRNA biogenesis necessarily relies on transcription initiating outside of the IS element and proceeding towards the *iscB* ORF **(Extended Data Fig. 1b)**. Our analyses therefore suggest that genomic insertions into transcriptionally active target sites aid in the generation of functional ωRNAs, and that these insertion products are either preferentially generated (during transposition) or preferentially retained. Notably, this orientation bias was absent for TnpB, whose ωRNA substrates rely on transcription that initiates within the IS element itself (described below), and for an unrelated *IS*630-family transposase that we included as a negative control (**Extended DataFig. 1c**).

To select TnpB and IscB candidates for experimental study, we sifted through our phylogenetic trees, prioritizing homologs that were found in the same species within related and high-copy TEs, indicative of recent transposition events. We converged on *Geobacillus stearothermophilus* (*Gst*), a thermophilic soil bacterium that has yielded useful thermostable proteins including Cas9^36^, reverse transcriptase^37^, and DNA polymerase^38^, and whose genome revealed a substantial expansion of five IS*605*-family elements encoding both TnpB and IscB, denoted IS*Gst1–5*, collectively comprising ∼1% of the entire genome (**Fig. 1c**)^39^. Analysis of small RNA sequencing data revealed that ωRNAs from multiple transposons were constitutively expressed (**Fig. 1d**)^36^, and we furthermore found that the left end (LE) and right end (RE) boundaries of these IS elements were highly similar in DNA sequence (**Extended Data Fig. 2a- d**), suggesting a common mechanism of mobilization. Using this information, we identified a candidate *tnpA* gene responsible for transposing these elements, as well as minimal non-autonomous IS elements that lacked protein-coding genes altogether and resembled palindrome-associated transposable elements (PATEs; **Extended Data Fig. 2e**)^40^.

**Figure 2.**
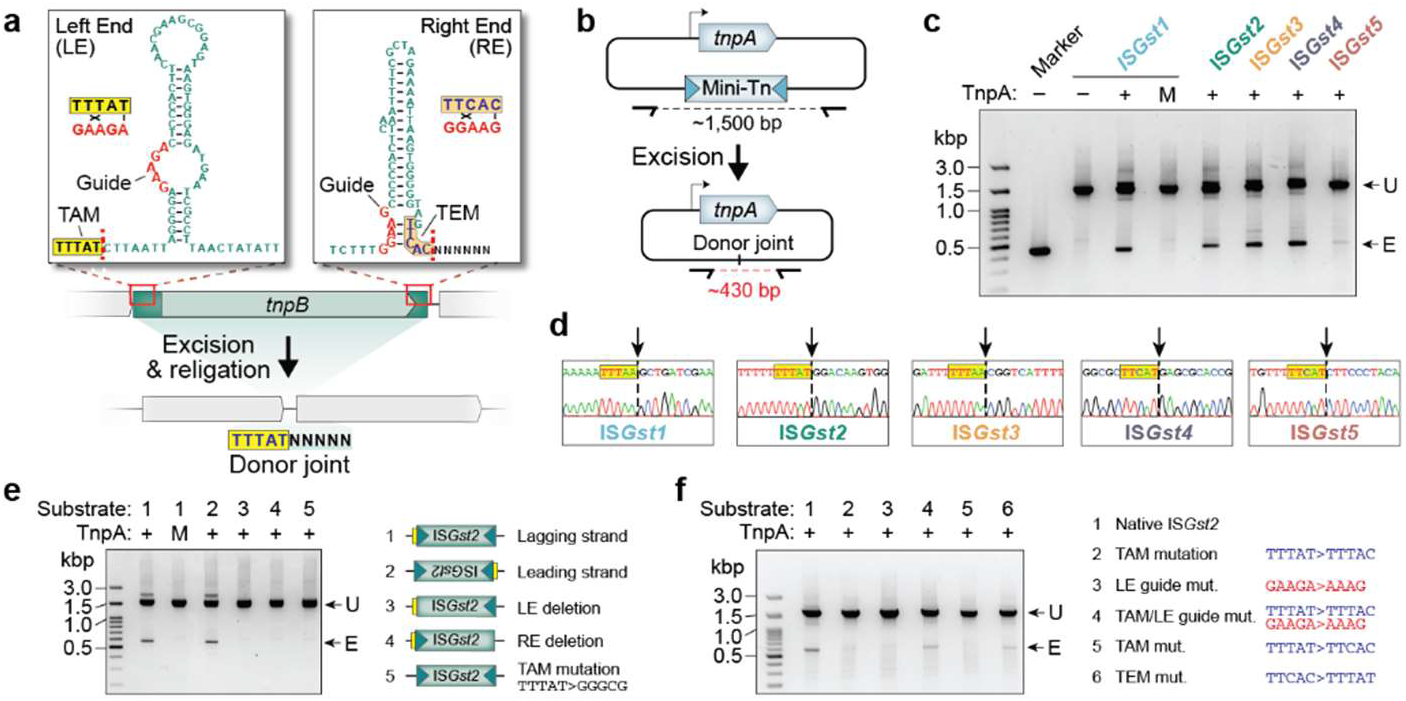
TnpA catalyzes DNA excision for multiple families of IS elements. **a,** Schematic of IS*Gst2* element, highlighting the subterminal palindromic transposon ends located on the top strand (top). Transposon-adjacent and transposon-encoded motifs (TAM and TEM) are highlighted in yellow and orange respectively, DNA guides are shown in red, and their putative base-pairing interactions are indicated; dotted lines indicate transposon boundaries and thus the sites of ssDNA cleavage and religation. The donor joint formed upon transposon loss is shown at the bottom and comprises the TAM abutting RE-flanking sequence (denoted with N’s). **b,** Schematic of heterologous transposon excision assay in *E. coli*. Plasmids encode TnpA and mini-transposon (Mini-Tn) substrates, whose loss is monitored by PCR using the indicated primers. **c,** TnpA is active in recognizing and excising all five families of IS*Gst* elements, as assessed by analytical PCR. Cell lysates were tested after overnight expression of TnpA with the indicated IS*Gst* mini-Tn substrates, and PCR products were resolved by agarose gel electrophoresis. Marker denotes a positive excision control; U, unexcised; E, excised; M denotes a Y125A TnpA mutant. **d,** Excision products from **c** exhibit the expected ‘donor joint’ architecture, as demonstrated by Sanger sequencing. Dotted lines denote the religation site following excision; the TAM is highlighted. **e,** Transposon excision requires intact LE and RE sequences, as seen via testing of the mutagenized mini-Tn substrates indicated on the right. Experiments were performed as in **c** using IS*Gst2*. Transposon ends and TAMs are indicated with green triangles and yellow boxes, respectively; M denotes a Y125A TnpA mutant. **f,** Transposon excision is dependent on cognate pairing between compatible TAM and guide sequences. Excision experiments were performed as in **c** using IS*Gst2* with the indicated mutations in the TAM/TEM (blue) or DNA guide (red). Substrate 4 has mutations to cognate sequences derived from IS*608*.

Interestingly, in addition to sharing similar sequences within the LE and RE, IS*Gst1–5* elements exhibited conserved, clade-specific transposon-adjacent motifs and transposon-encoded motifs (TAMs and TEMs; **Extended Data Fig. 2a-d**). Prior studies on the TnpA transposase from *Helicobacter pylori* IS*608*, which transposes a related IS*605*-like element, revealed that these motifs constitute the target and cleavage sites recognized during transposon insertion and transposon excision reactions, respectively. Yet rather than being recognized exclusively through protein-DNA recognition, these motifs form non-canonical base-pairing interactions with a DNA ‘guide’ sequence located in the sub-terminal ends of the IS element (**Fig. 2a**)^41,42^. Focusing on multiple sequence alignments between IS*Gst1–5* elements, we observed covarying mutations between both the TAM/TEM sequences and their associated DNA guide sequences (**Fig. 2a, Extended Data Fig. 2a,b**), further suggesting that these elements would be active for transposition.

### *Gst*TnpA is active for DNA excision and transposition

TnpA is part of the HUH endonuclease superfamily and recognizes DNA hairpin structures in a sequence and structure-specific way to mediate ssDNA cleavage and ligation^22^. Biochemical and genetic studies of an IS*608*-encoded TnpA homolog from *H. pylori* (*Hpy*TnpA) revealed a peel-out-paste-in transposition mechanism, in which IS elements first excise as a circular single-stranded DNA (ssDNA) intermediate with abutted LE and RE sequences, followed by precise rejoining of the flanking DNA to produce a scarless ‘donor joint’ that regenerates the original genomic sequence^23^. Transposition to a new target site occurs downstream of a TAM recognized through base-pairing with the LE DNA guide, via insertion of the circular ssDNA in a manner that requires subsequent second-strand synthesis but does not create a target-site duplication. Excision and integration reactions both rely heavily on ssDNA and the formation of conserved, intramolecular LE and RE stem-loop structures (**Fig. 2a**), implicating DNA replication as a major opportunity for transposon activation^23,41^.

We designed a DNA excision assay to test the activity of *Gst*TnpA on a mini-transposon (mini-Tn) substrate derived from its native autonomous IS element, IS*Gst1*. We cloned *E. coli* expression vectors that encoded *Gst*TnpA upstream of the mini-Tn, which comprised an antibiotic resistance gene flanked by full-length LE and RE sequences and genomic *G. stearothermophilus* sequences upstream and downstream of the predicted transposon boundaries. Primers were designed to bind outside the mini-Tn, such that PCR from cellular lysates would amplify either the starting substrate or a shorter reaction product resulting from transposon excision and religation (**Fig. 2a,b**). We also generated a parallel panel of substrates containing LE and RE sequences derived from IS*Gst2–5*, which natively encode IscB, TnpB, or ωRNA only, to determine the breadth of *Gst*TnpA substrate recognition. Remarkably, *Gst*TnpA was active on all five families of IS elements, with excision dependent on the predicted catalytic tyrosine residue (**Fig. 2c)**, but failed to cross-react with a DNA substrate derived from the *H. pylori* IS*608* element (**Extended Data Fig. 3a,b**). Sanger sequencing of excision products revealed that in each case, TnpA precisely re-joined sequences flanking the mini-Tn to generate a scarless donor joint (**Fig. 2d**), which we hypothesized could be recognized and cleaved by TnpB/IscB (see below). Using an alternative qPCR-based strategy to prime directly off the donor joint sequence, we calculated excision frequencies of 0.70% directly from overnight cultures (**Extended Data Fig. 3c-e**).

**Figure 3.**
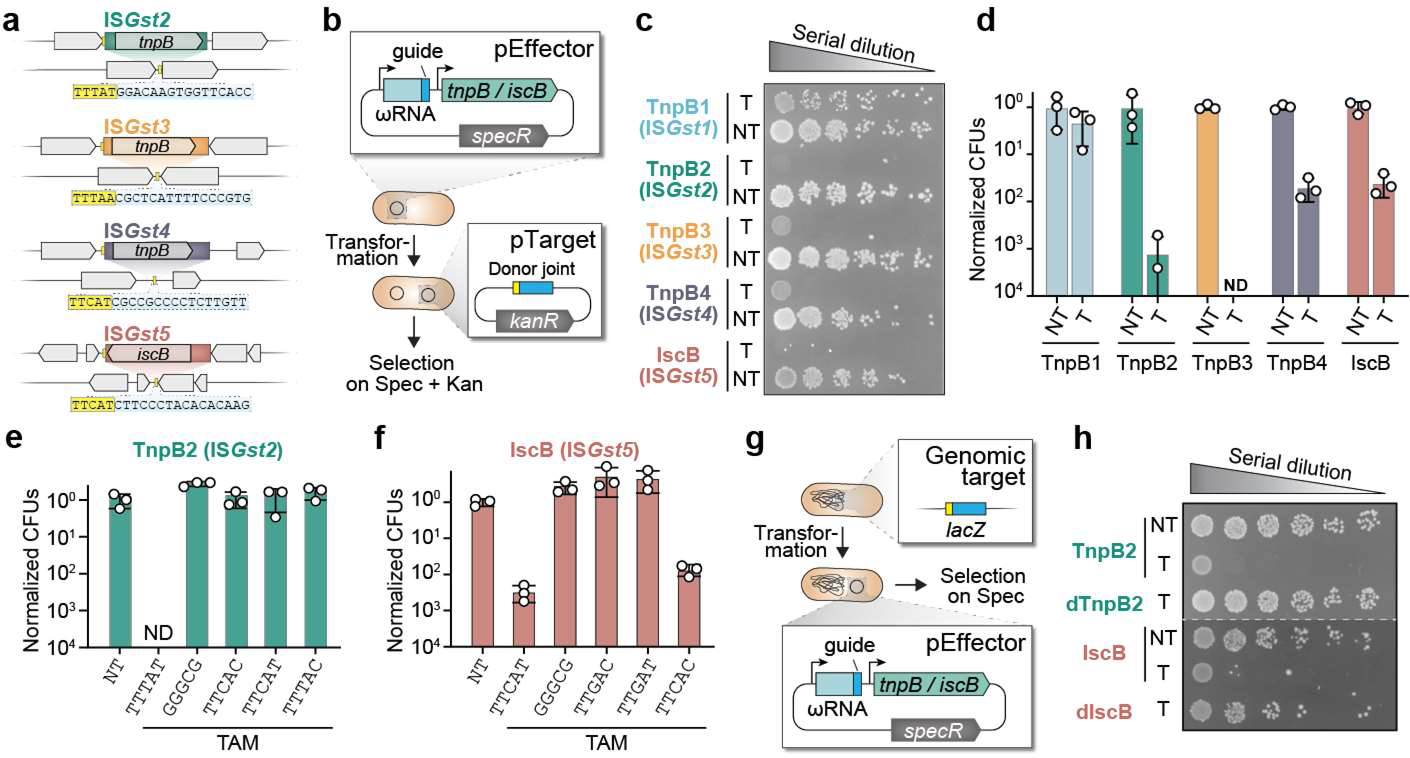
TnpB and IscB target ‘donor joint’ molecules excised by TnpA. **a,** Schematic representation of each IS family (colored rectangle), alongside homologous sites from related *Gst* strains that lack the transposon insertion. TAMs are highlighted in the donor joint sequences shown below each element. **b,** Schematic of *E. coli*-based plasmid interference assay. Protein-RNA complexes are encoded by pEffector, and targeted cleavage of pTarget results in a loss of kanamycin resistance and cell lethality on selective LB-agar plates. **c,** *G.stearothermophilus* TnpB and IscB homologs are highly active for RNA-guided DNA cleavage, as assessed by plasmid interference assays. Transformants with a targeting (T) or non-targeting (NT) ωRNA-pTarget combination were serially diluted and plated on selective media at 37 °C for 24 h. **d,** Quantification of the data in **c**, normalized to the non-targeting (NT) plasmid control for each IS*Gst* element. CFU, colony forming units; ND, not detected. **e,** DNA cleavage by TnpB2 is highly sensitive to TAM mutations, as assessed by plasmid interference assays. Data were quantified and plotted as in **d** for the indicated TAM mutations; TTTAT denotes the WT TAM. **f,** DNA cleavage by IscB is highly sensitive to TAM mutations, as assessed by plasmid interference assays. Data were quantified and plotted as in **d** for the indicated TAM mutations; TTCAT denotes the WT TAM. **g,** Schematic of *E. coli*-based genome targeting assay, in which RNA-guided DNA cleavage of *lacZ* by TnpB/IscB results in cell death. **h,** TnpB2 and IscB are active for targeted genomic DNA cleavage, as assessed by genome targeting assay. Transformants with a targeting (T) or non-targeting (NT) ωRNA were serially diluted and plated on selective media at 37 °C for 24 h. dTnpB2, D196A mutation; dIscB, D58A/H209A/H210A mutations.

We next investigated sequence determinants of transposon excision in greater detail, focusing on the IS*Gst2* element that natively encodes TnpB and its associated ωRNA. Excision proceeded regardless of whether the mini-Tn was encoded on the leading or lagging-strand template, but was ablated when we scrambled either the LE or RE sequence, confirming the critical importance of these regions for TnpA recognition. Excision was also strongly dependent on the presence of a cognate TAM adjacent to the LE as well as a compatible DNA ‘guide’ sequence located within the LE, since mutation of either region led to a loss of product formation (**Fig. 2e, S2a**). Interestingly, however, simultaneous mutation of both the TAM and LE guide sequence to the corresponding motifs found in IS*608* restored excision activity with *Gst*TnpA (**Fig. 2f**), confirming the importance of these complementary base-pairing interactions to mediate site-specific DNA excision. Similar base-pairing interactions occur between a DNA ‘guide’ sequence within the RE and a matching TEM found at the RE boundary, with only minor differences between the TAM and TEM at positions 3 and 5 (**Extended Data Fig. 2a,b**). Whereas the excision reaction did not tolerate mutation of the TAM sequence to the TEM sequence, we were surprised to find that mutations to the TEM were still tolerated, despite ablating predicted base-pairing interactions with the RE ‘guide’ sequence (**Fig. 2f**). However, closer inspection revealed that these excision events resulted from erroneous selection of an alternative mini-Tn boundary downstream of the native RE, at a sequence matching the WT TEM (TTCAC; **Extended Data Fig. 3f,g**). These results indicate that IS*200*/IS*605*-family elements tolerate flexible spacing between the TAM/TEM and corresponding guide sequences, potentially allowing for capture of additional sequences outside of the native LE and RE boundaries.

We also investigated the ability of *Gst*TnpA to catalyze transposon insertions at new target sites. Using a traditional mating-out assay with the IS*Gst2* mini-Tn (**Extended Data Fig. 4a**), in which transposition events into a conjugative plasmid are isolated via drug selection, we measured transposition efficiencies of 2.5 x 10^-7^, which were several orders of magnitude lower than the observed rates of transposon excision (**Extended Data Fig. 3e,4b**). These results suggest that, under the tested experimental conditions, TnpA expression would eventually lead to permanent transposon loss from the cell population, absent any active mechanisms for maintaining transposons at their donor sites during or after excision (see below). Long-read sequencing of drug-resistant transconjugants confirmed the presence of novel mini-Tn insertions, which were invariably located downstream of endogenous TAM sites on the F-plasmid, confirming the essentiality of this motif for both mini-Tn excision and integration (**Extended Data Fig. 4c**). Collectively, these experiments demonstrate that *Gst*TnpA is active in mobilizing a large network of diverse, IS*605*-like elements found in the *G. stearothermophilus* genome, but that its intrinsic enzymatic properties render transposons vulnerable to being permanently lost from the population without an active mechanism for donor-site preservation. Thus, we next focused on the molecular properties of IscB and TnpB, given their frequent presence as accessory factors encoded within the very same transposons.

**Figure 4.**
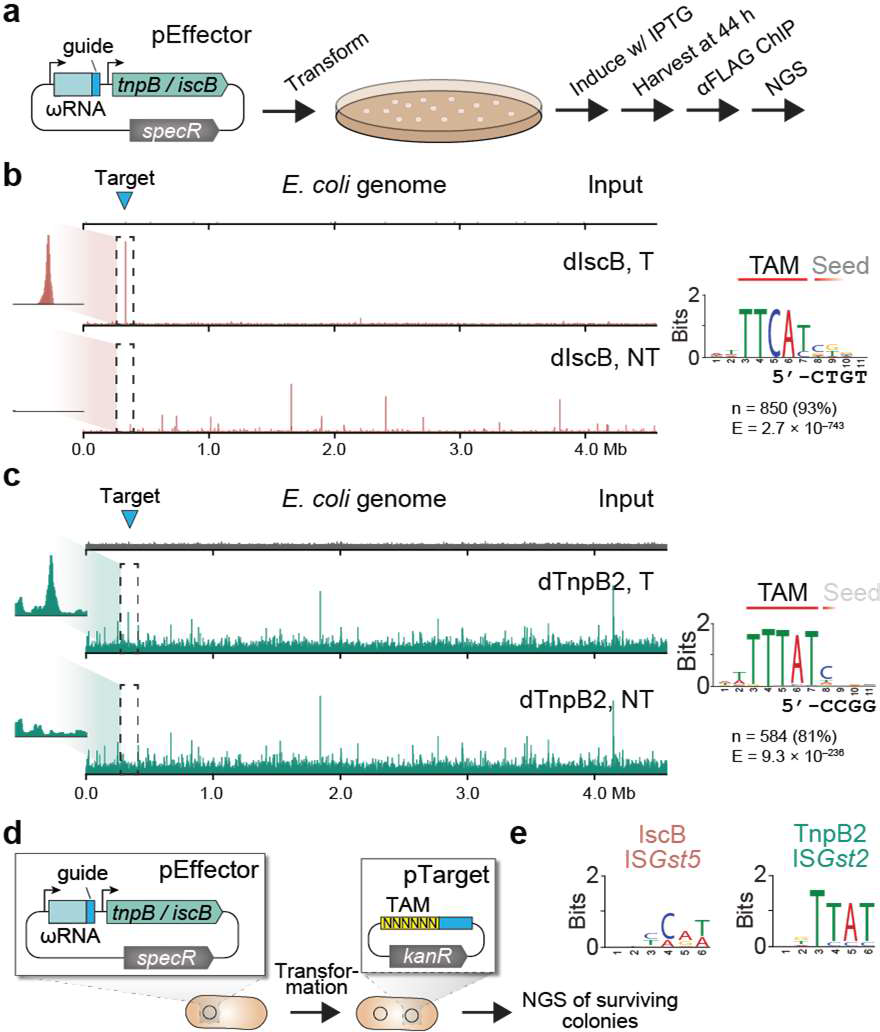
Unbiased identification of TnpB/IscB TAM specificity by ChIP-seq and library assays. **a,** Schematic of ChIP–seq workflow to monitor genome-wide binding specificity of TnpB/IscB. *E. coli* cells were transformed with plasmids encoding catalytically inactive dTnpB2 or dIscB and a genome targeting (T) or non-targeting (NT) ωRNA. After induction, cells were harvested, protein–DNA cross-links were immunoprecipitated, and NGS libraries were prepared and sequenced. **b,** (Left) Genome-wide representation of ChIP-seq data for dIscB with target site (blue triangle) shown, for T and NT samples alongside the input control. Coverage is shown as reads per kilobase per million mapped reads (RPKM), normalized to the highest peak in the T sample. (Right) Off-target binding events were analyzed by MEME ChIP, which revealed a strongly conserved consensus motif consistent with the WT TAM (TTCAT) but weak seed sequence bias; part of the ωRNA guide sequence is shown below. Consensus motifs are oriented 5’ of the IS element left end. n, number of peaks contributing to the motif; E, E-value significance. **c,** Representative ChIP-seq data for dTnpB2, plotted as in **b**. **d,** Schematic of TAM library cleavage assay, in which plasmids expressing nuclease-active TnpB/IscB and an associated ωRNA (pEffector) are designed to cleave a target sequence flanked by randomized 6-mer (pTarget). Plasmid cleavage results in plasmid elimination, loss of cell viability, and depletion of the particular TAM upon library sequencing. **e,** WebLogo representation of the 10-most depleted sequences upon deep sequencing of plasmid samples from the TAM library cleavage assay for TnpB2 and IscB Consensus motifs are oriented 5’ of the IS element left end.

### *Gst*TnpB and IscB homologs function as RNA-guided endonucleases

We hypothesized that TnpB/IscB nucleases might function with transposon-encoded ωRNAs to target the ‘donor joint’ produced upon scarless transposon excision (**Fig. 3a**), thereby forcing cells to survive otherwise lethal DNA DSBs by restoring another copy of the transposon through recombination^27,43,44^. With knowledge that *Gst*TnpA was active in mobilizing diverse IS elements, we turned our attention to reconstituting nuclease activity for the associated *Gst*TnpB/IscB proteins using a plasmid interference assay, in which successful targeting leads to plasmid cleavage and a loss of cellular viability (**Fig. 3b**). We designed expression plasmids encoding both TnpB/IscB and the corresponding ωRNA guides derived from their native *Gst*IS elements (pEffector), alongside target plasmids containing donor joints that were bioinformatically identified and experimentally verified in TnpA excision assays (pTarget; **Fig. 2d,3a**). After screening various promoter combinations driving expression of the nuclease and ωRNA (**Extended Data Fig. 5a**), we found that *Gst*IscB and three *Gst*TnpB distinct homologs were highly active for RNA-guided DNA cleavage of their native donor joints (**Fig. 3c,d**). Interestingly, *Hpy*TnpB encoded by the well-studied IS*608* element was inactive when tested under similar conditions, whereas we recapitulated the recently described activity for *Dra*TnpB (**Extended Data Fig. 5b**)^27^.

**Figure 5.**
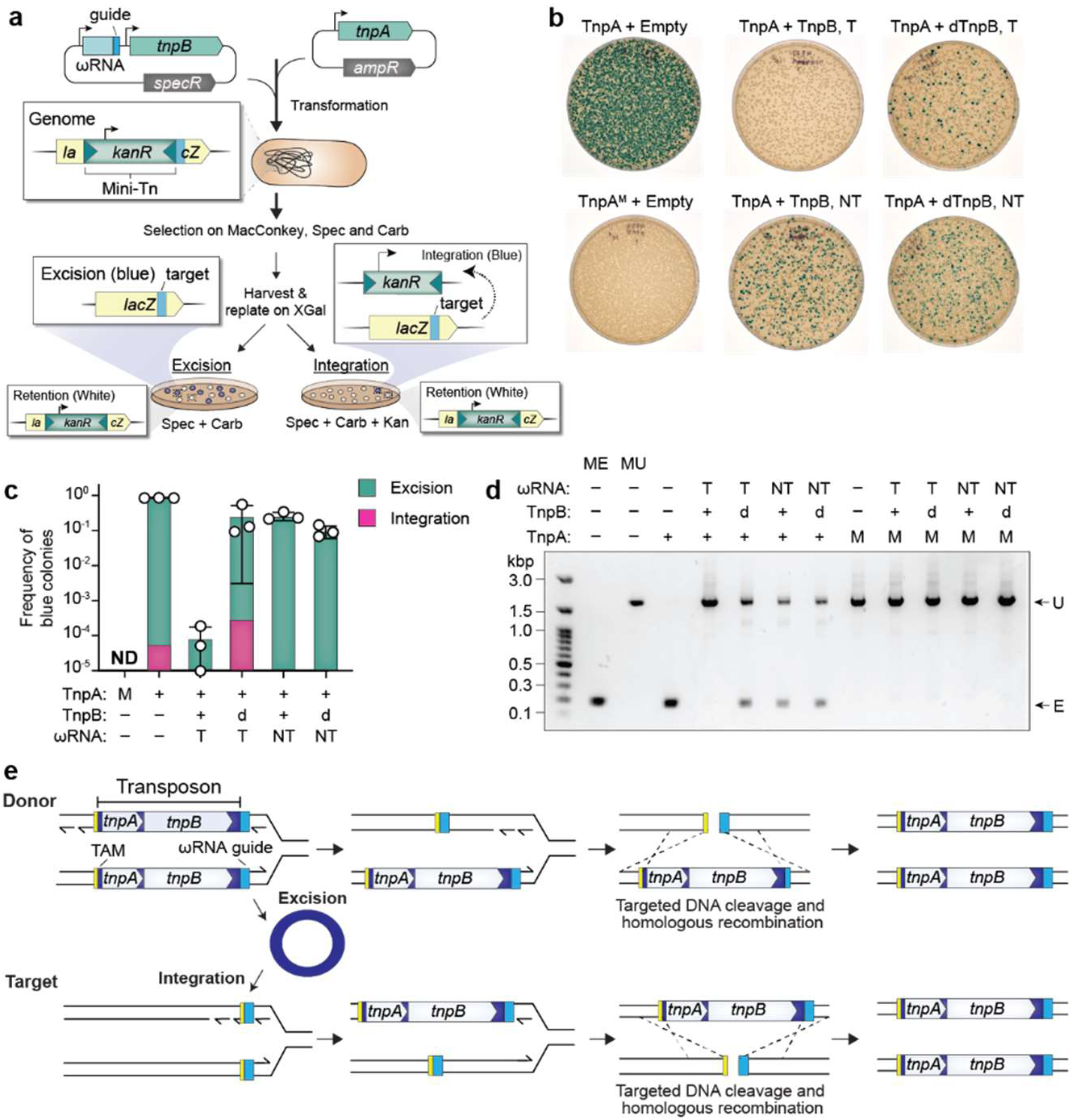
RNA-guided nucleases preserve IS elements at the donor site following transposase-mediated excision. **a,** Schematic of experimental workflow to measure transposon fate in *E. coli* in the presence of TnpA and TnpB. A mini-Tn was inserted at a compatible TAM site in *lacZ*, such that cells grown on X-gal exhibit a blue colony phenotype upon permanent transposon excision, or a white colony phenotype if the transposon is retained. Cells were transformed with plasmids expressing WT or mutant TnpA and/or TnpB2, with a targeting (T) or non-targeting (NT) ωRNA. Kanamycin-resistant cells with a blue color phenotype will result upon transposon loss at the donor site (excision) and transposon gain at a new target site (integration). **b,** TnpB promotes robust transposon retention at the donor site, as assessed by blue-white colony screening. Representative plating results are shown from experiments that included the indicated components. M, TnpA (Y125A) mutant; dTnpB, D196A mutation. **c,** Quantification of the data from **b** across multiple experimental replicates. Green bars indicate the frequency of blue colonies as a measure of transposon excision/retention for the indicated experimental conditions and pink indicates the frequency of blue colonies that maintain kanamycin resistance as a measure of transposon excision and reintegration elsewhere in the genome. Bars indicate mean ± standard deviation (n = 3). **d,** Genotypes inferred from blue-white colony screening were assessed by PCR analysis and agarose gel electrophoresis for the indicated experimental conditions, which reports on whether the mini-Tn is unexcised (UE) or excised (E) at the donor *lacZ* site. The first two lanes denote marker controls (ME: Mock excised and MU: Mock Unexcised) for the two possible PCR products. **e,** Peel-and-paste/cut-and-copy model for how TnpA and TnpB/IscB coordinate their catalytic activities to maintain the presence of IS*200/605*-family transposons at donor sites. TnpA mediates excision and religation of flanking sequences at the donor site as ssDNA becomes available during DNA replication, resulting in transposon loss from the donor site. The excised ssDNA product is concurrently ligated to form a circular ssDNA-transposome complex, which can be reintegrated downstream of a TAM motif elsewhere in the genome, albeit at much lower efficiency than excision. In the presence of TnpB/IscB, RNA-guided DNA cleavage of the donor joint initiates homologous recombination with the sister chromosome that still contains the IS element, thus rapidly restoring the transposon at the original donor site; the absence of TnpB/IscB leads to permanent transposon loss after cell division. TnpB/IscB can also cleave sister chromosomes lacking the newly integrated IS element after transposition to a new target site, facilitating further spread. The transposon is shown in dark blue; the TAM is shown in yellow, and light blue rectangles represent regions complementary to the guide portion of the ωRNA.

An elegant feature of IS*605*-family elements is that the same LE-abutting TAM sequence that TnpA requires for transposon excision and integration is also presumably required by TnpB/IscB for DNA targeting and cleavage, akin to the role of PAM sequences with CRISPR- Cas9 and Cas12^26,27,45^. We systematically mutagenized the TAM on pTarget and found that DNA cleavage was ablated with even single-bp changes, which importantly, would also render the site of ωRNA biogenesis at the transposon RE — where the motif differs from the cognate TAM in only two positions — completely unrecognizable (**Fig. 3e,f and** **Extended Data Fig. 5c,d**). TnpB and IscB were both functional for genomic targeting and cleavage as well, and point mutations in the predicted HNH and/or RuvC nuclease domains completely ablated activity (**Fig. 3g,h**). Interestingly, a panel of three TnpB-specific ωRNAs targeting *lacZ* showed varying levels of activity, as assessed by cell lethality (**Extended Data Fig. 5e**), suggesting additional unknown requirements that impact the efficiency of DNA targeting and cleavage.

To investigate binding specificity in more detail, we next performed ChIP-seq experiments to map all chromosomal binding sites of nuclease-dead IscB and TnpB programmed with *lacZ*- specific ωRNAs (**Fig. 4a**). The resulting data revealed strong enrichment at the on-target site and numerous off-targets (**Extended Data Fig. 6a-d**), and the majority of peaks shared highly conserved consensus motifs of 5’-TTCAT-3’ (IscB from IS*Gst5*) and 5’-TTTAT-3’ (TnpB2 from IS*Gst2*) (**Fig. 4b,c**), which precisely matched the TAM motifs neighboring the native IS*Gst5* and IS*Gst2* elements, respectively (**Extended Data Fig. 2a**). Similar consensus motifs emerged when we tested cleavage activity in cells using pTarget libraries containing degenerate TAM sequences (**Fig. 4d**), indicating common sequence determinants for DNA target binding and cleavage. Interestingly, neither IscB nor TnpB exhibited a strong requirement for extensive complementarity within the seed sequence for the off-target sites analyzed (**Extended Data Fig. 6a,b**), and this absence was particularly striking in comparison to matched experiments with Cas9 and Cas12a, which were strongly dependent on 3–5 nt of PAM-adjacent sequence matching the guide RNA (**Extended Data Fig. 6c,d**)^46^. These results suggest the possibility that Cas9 and Cas12 may have evolved a greater degree of reliance on RNA-DNA complementarity for stable DNA binding, whereas IscB and TnpB may be strictly dependent on a more extensive TAM motif but permissive of RNA-DNA mismatches^47,48^.

Taken together with data highlighting the importance of the TAM in TnpA-mediated transposon excision and integration, these findings implicate the TAM as a critical hub of DNA interrogation during multiple stages of the transposon life cycle for IS*200*/IS*605*-family elements. Despite the chemically diverse reactions being catalyzed during DNA transposition and DNA cleavage, IscB/TnpB nucleases and TnpA transposases were both constrained by selective pressures to faithfully recognize overlapping sequence motifs, presumably for selfish interests to promote transposon proliferation. We therefore focused our subsequent efforts on investigating how TnpA and TnpB coordinate their biochemical activities to modulate transposon lifestyle when expressed simultaneously in a heterologous system.

### RNA-guided nucleases promote transposon retention through targeted DSBs

We hypothesized that co-expression of IscB/TnpB nucleases with compatible ωRNAs would rapidly intercept the donor joint products generated upon transposon excision by TnpA, thereby creating a targeted DNA double-strand break (DSB) at the site of excision. DSBs are typically lethal in bacteria unless they can be acted upon and repaired by homologous recombination^43,44^, and the strand-specific excision of IS*200*/IS*605*-family transposons as ssDNA would ensure that the sister chromosome produced during DNA replication can always serve as a homologous donor template. Thus, the mechanism we envisioned would be akin to the role of homing endonucleases in promoting lateral mobility through DSB-triggered recombination^49^, but rather than specifying new target sites for transposon insertion, IscB/TnpB would promote reinstallation of transposon copies at pre-existing donor sites.

To test this, we generated an *E. coli* strain harboring a *lacZ*-interrupting mini-Tn that was inserted downstream of a TnpB-compatible TAM, such that scarless excision by TnpA would result in a phenotypic switch from *lacZ^−^* (white colony phenotype) to functional *lacZ^+^* (blue colony phenotype; **Fig. 5a**). We transformed strains with expression plasmids encoding TnpA (or an inactive mutant) and TnpB (or an inactive mutant), programmed with either a non-targeting ωRNA or a *lacZ*-targeting ωRNA designed to cleave the donor joint generated upon TnpA-mediated mini-Tn excision. After enriching for excision events by growing strains on MacConkey agar, we plated cells on media containing X-gal and performed blue-white colony screening. Using this approach, we immediately observed the emergence of a large fraction of blue colonies in the presence of WT TnpA, but not a catalytically inactive mutant, and colony PCR analysis confirmed that these colonies had indeed permanently lost the transposon at the donor *lacZ* locus (**Fig. 5b-d**). When we plated a similar population of cells onto X-gal plates that also contained kanamycin, thus selecting for the presence of the mini-Tn, blue colonies were 1000X less abundant (**Fig. 5c**), confirming our earlier findings that the frequency of transposon excision at the donor site vastly exceeds the frequency of transposon integration at a new target site.

Remarkably, co-expression of TnpB and a *lacZ*-specific ωRNA completely eliminated the emergence of blue colonies under otherwise identical conditions, and colony PCR confirmed that transposons were uniformly maintained at their original genomic location (**Fig. 5b-d, Extended Data Fig. 7**). Importantly, this phenotypic effect was dependent on both a targeting ωRNA and an intact TnpB nuclease domain, indicating that targeting/binding alone is insufficient for transposon retention at the donor site, but that targeted cleavage and local DSB formation are necessary. These results highlight the importance of TnpB nucleases to faithfully and robustly preserve transposons at the donor site that are otherwise lost via TnpA-mediated excision, through formation of targeted DSBs and ensuing recombination (**Fig. 5e**).

## DISCUSSION

Prokaryotic insertion sequences are among the simplest type of transposable element found in nature, in that they are often defined as minimal elements that contain only those gene(s) required for transposition. Paradoxically, however, large families of highly abundant IS elements carry genes that do not encode active transposases, but rather, encode diverse accessory genes that provide myriad selective advantages for survival of the transposable element, including resolution systems, antibiotic resistance genes, and pathogenicity functions^40^. In the case of IS*200*/IS*605*- family elements, the presence of pervasive accessory genes accompanying *tnpA* has long been observed and defined with variable nomenclatures (*tnpB, orfB, tlpB*, etc.), but aside from experiments demonstrating that these gene products are not required for transposition^24,25,32,50^, the molecular function of TnpB-family proteins remained mysterious. Two recent studies provided critical new insights into the biochemical activities of both TnpB and a related accessory protein, IscB, by demonstrating that both enzymes employ transposon-encoded guide RNAs, termed ωRNAs, to catalyze RNA-guided DNA cleavage. Yet despite shedding light on the intriguing evolutionary origins of CRISPR-Cas9 and Cas12 effector nucleases, these studies did not explore the role that TnpB/IscB play in the context of transposition^20,26,27^.

Our results reveal that RNA-guided DNA endonucleases encoded within IS*200*/IS*605*- family transposons evolved a “cut-and-copy” mechanism to direct site-specific DNA double-stranded breaks and homologous recombination, to ensure the retention and proliferative spread of these elements. The fact that these elements couple ssDNA transposition to DNA replication, given the enzymatic mechanism of Y1 TnpA transposase^51^, elegantly guarantees that a donor template is always available for DSB repair following strand-specific DNA transposition, and the resulting scarless excision and integration products furthermore ensure faithful homology between both molecules during recombination. We propose that this pathway is not only essential to retain the transposon at the original donor site, but that it also drives the element to spread to sister chromosomes, likely increasing the transposon copy number over many generations as new donor substrates are available to be mobilized in *trans* by TnpA. Indeed, the genome of *G. stearothermophilus* contains nearly 50 elements that lack *tnpA* but that we predict could be mobilized *in trans*, based on sequence conservation of the transposon ends. Similar observations have been made in other bacteria, suggesting that autonomous elements may be unstable or prone to degradation^20^. The very presence of PATE elements, which often encode ωRNAs that would themselves need to be acted upon *in trans* by TnpB proteins encoded within distinct IS elements, supports the modularity of these molecular components^26,40^.

Class II (DNA) transposons have evolved diverse mechanisms of mobilization, which differ critically in the fate of DNA at the donor site, the form of the mobile DNA itself, and the chemical mechanism by which the mobile DNA is integrated at the new target site. Conventional cut-and-paste transposons are excised as dsDNA molecules and integrated in a replication-independent fashion, leaving behind DSBs at the donor site that can be acted upon in a variety of ways^52^. Copy-and-paste transposons depend on local replication for transposition, and as such, are always retained at their original donor site after integration occurs at the new target site. Last but not least, peel-out-paste-in transposons like IS*200*/IS*605* excise as ssDNA molecules from only one strand of the donor molecule, concomitant with rejoining of the flanking sequences to create a scarless donor joint^53^. Despite these distinct lifestyles, evolutionary survival of the selfish element ultimately requires that the rate of transposon loss – whether by permanent excision, inactivation, or silencing – be compensated by transposon gain, through novel integration events. Our experiments demonstrate the TnpA enzymes catalyze transposon excision (i.e. loss) at rates that exceed transposon integration (i.e. gain) by orders of magnitude, which would be seemingly incompatible with the long-term survival of these elements. Yet the powerful combination of TnpA and TnpB/IscB-family nucleases, which evolved to use localized guide RNAs to force recombination at the sites of excision and integration, offers a unique solution to this problem by achieving TnpA-mediated ‘peel-and-paste’ transposition, coupled with TnpB/IscB-mediated ‘cut-and-copy’ super-Mendelian inheritance, through the sequential steps of excision, cleavage, and recombination (**Fig. 5e**).

There are interesting parallels between the transposon retention mechanism by TnpB-family nucleases, and the mobilization mechanism of group I introns that encode site-specific homing endonucleases (HEs). In that case, HE enzymes target conserved sites for DSB formation, thereby promoting a type of transposition that relies entirely on recombination to copy the element from the donor to the target. IS*200*/IS*605*-family elements have instead harnessed programmable nucleases as ‘adaptive’ site-specific enzymes that target the empty donor site programmed after each transposition event, to selfishly bias the inheritance and retention of the transposable element for its continued spread. Intriguingly, IS*200*/IS*605*-family elements themselves are sometimes encompassed within group I introns^50^, highlighting the diversity of these elements and their associated gene products.

The combination of DNA-guided transposition (TnpA) and RNA-guided nuclease activity (TnpB/IscB) activity has facilitated the pervasive spread of IS*200*/IS*605*-family elements throughout bacteria and archaea. Remarkably, however, TnpB homologs have also been identified within diverse transposable elements and in association with distinct transposases that mobilize DNA through a variety of different mechanisms. Furthermore, these homologs have been identified in all domains of life, including higher eukaryotes, suggesting that the very biochemical property of RNA-guided DNA targeting and cleavage is not only restricted to CRISPR-associated enzymes (Cas9 and Cas12) and IS element-associated enzymes (IscB and TnpB), but to nucleases that are broadly present throughout life^28-30^. In this regard, CRISPR-Cas systems represent just one powerful example whereby RNA-guided DNA targeting nucleases within the TnpB superfamily have been exapted and repurposed for antiviral defense. The diverse and widespread phylogenetic distribution of these enzymes strongly suggest that other cellular pathways will be identified where RNA-guided targeting enzymes were similarly coopted for new molecular functions.

## METHODS

### Data reporting

No statistical methods were used to predetermine sample size. The experiments were not randomized, and investigators were not blinded to allocation during experiments and outcome assessment.

### IscB and TnpB detection and database curation

Homologs of IscB proteins were comprehensively detected using the amino acid sequence of a *K. racemifer* homolog (NCBI Accession: WP_007919374.1) as the seed query in a JackHMMER part of the HMMER suite (v3.3.2). To minimize false homologs, a conservative inclusion and reporting threshold of 1e-30 was used in the iterative search against the NCBI NR database (retrieved on 06/11/2021), resulting in 5,715 hits after convergence. These putative homologs were then annotated to profiles of known protein domains from the Pfam database (retrieved on 06/29/2021) using hmmscan with an E-value threshold of 1e-5. Proteins that did not contain the RRXRR, RuvC, RuvC_III, or the RuvX domain were discarded. Although the HNH domain was annotated, proteins without the HNH were not removed. The variation in the presence of the HNH domain was preserved to better represent the natural diversity of IscBs. From the remaining set, proteins that were less 250 aa were removed to eliminate partial or fragmented sequences, resulting in a database of 4,674 non-redundant IscB homologs. Contigs of all putative *iscB* loci were retrieved from NCBI for downstream analysis using the Bio.Entrez package.

TnpB homologs were comprehensively detected similarly to IscB, use both the *H. pylori* (*Hpy*TnpB) amino acid sequence (NCBI Accession: WP_078217163.1) and the *G. stearothermophilus* (*Gst*TnpB2) amino acid sequence (NCBI Accession: WP_047817673.1) as seed queries for two independent iterative jackhammer searches against the NR database, with an inclusion and reporting threshold of 1e-30. The union of the two searches were taken, and proteins that were less than 250 aa were removed to trim partial or fragmented sequences, resulting in a database of 95,731 non-redundant TnpB homologs. Contigs of all putative *tnpB* loci were retrieved from NCBI for downstream analysis using the Bio.Entrez package.

### Phylogenetic analyses

IscB protein sequences were clustered with at least 95% length coverage and 95% alignment coverage using CD-HIT^54^ (v4.8.1). The clustered representatives were taken and aligned using MAFFT^55^ (v7.508) with the E-INS-I method for 4 rounds. Post-alignment cleaning consisted of using trimAl^56^ (v1.4.rev15) to remove columns containing more than 90% of gaps and manual inspection. The phylogenetic tree was created using IQ-Tree 2^57^ (v2.1.4) with the WAG model of substitution. Branch support was evaluated with 1000 replicates of SH-aLRT, aBayes, and ultrafast bootstrap support from the IQTREE package. The tree with the highest maximum-likelihood was used as the reconstruction of the IscB phylogeny.

Putative TnpB sequences were clustered by 50% length coverage and 50% alignment coverage using CD-HIT^54^. Similar to IscB, the clustered representatives were taken and aligned using MAFFT^55^ with the E-INS-I method for 4 rounds. Post-alignment cleaning consisted of using trimAl^56^ to remove columns containing more than 90% of gaps and manual inspection. The phylogenetic tree was created using IQ-Tree 2^57^ with the WAG model of substitution. Branch support was evaluated with 1000 replicates of SH-aLRT, aBayes, and ultrafast bootstrap support from the IQTREE package. The tree with the highest maximum-likelihood was used as the reconstruction of the TnpB phylogeny.

### ωRNA covariation analyses

Initially searches of the Rfam database indicated a potential ncRNA belonging to the HNH endonuclease-associated RNA and ORF (HEARO) RNA (RF02033)^33,58^. A covariance model of HEARO RNA (retrieved 6/24/2021) was initially used to discover all HEAROs within our curated IscB-associated contig database using cmsearch from the Infernal package (v1.1.4)^59^. A liberal minimum bit score of 15 was used in an attempt to capture distant or degraded HEAROs, and the identification of a HEARO as a putative ωRNA was supported by its proximity, orientation, and relative location to the nearest identified IscB ORF. Remaining hits were considered ωRNAs if they were upstream of an IscB ORF and within 500 bp or overlapping with the nearest IscB ORF. After inspecting the RF02033 model, it appeared to lack additional structural elements located downstream. To address this, the boundaries of ωRNA were refined and used to generate a more accurate, comprehensive covariance model. Hits to the RF02033 model described above were retrieved, expanded 200 bp downstream, and clustered by 80% length coverage and 80% alignment coverage using CD-HIT^54^. CMfinder^60^ (v0.4.1.9) was then used with recommended parameters to discover new motifs *de novo*. Additional structures were discovered and present in over 80% of the expanded sequences. This covariance model was used to expand the 3’ coordinates of previously identified ωRNAs to encompass the second stem loop using cmsearch^59^ on expanded ωRNAs. These refined ωRNA boundaries and sequences were then used to create a new ωRNA model. The refined ωRNAs were clustered by 99% length coverage and 99% alignment coverage using CD-HIT^54^ to remove duplicates. A structure-based multiple alignment was then performed using mLocARNA^61^ (v1.9.1) with the following parameters:

> *--max-diff-am 25 --max-diff 60 --min-prob 0.01 --indel -50 --indel-open -750 --plfold-span 100 --alifold-consensus-dp*

The resulting alignment with structural information was used to generate a new ωRNA covariance model with the Infernal suite, refined with Expectation-Maximization from CMfinder^60^, and verified with R-scape at an E-value threshold of 1e-5. The resulting ωRNA covariance model was used with cmsearch^59^ to discover new ωRNAs within our curated IscB-associated contig database. The resulting sequences were aligned to generate a new CM model that was used to again search our IscB-associated contig database. This process was repeated three times for our final generic IscB-associated ωRNA model.

While covariance models of TnpB-associated ωRNAs were available through Rfam (RF03065) and (RF02998), these models appeared to only include a very small subset of TnpB-associated ωRNA and contained very few hits. Based on small RNA-seq analysis that suggested a ncRNA often overlapped with the TnpB ORF and extending into the RE boundary of the IS element, we extracted sequences 150 bp downstream of the last nucleotide of the TnpB ORF to define the RE and transposon boundaries. The ∼150-bp sequences were clustered by 99% length coverage and 99% alignment coverage using CD-HIT^54^ to remove duplicates. The remaining sequences were then clustered again by 95% length coverage and 95% alignment coverage using CD-HIT^54^. This was done to identify clusters of sequences that were closely related but not identical, as expected of IS elements that have recently mobilized to new locations. For the 300 largest clusters, which all had a minimum of 10 sequences, MUSCLE^62^ (v3.8.1551) with default parameters was used to align each cluster of sequences. Then, each cluster alignment was manually inspected for the boundary between high conservation and low conservation, or where there was a stark drop-off in mean pairwise identity over all sequences. This point was annotated for each cluster as the putative 3’ end of the IS elements. If there was no conservation boundary, sequences in these clusters were expanded by another 150 bp, in order to capture the transposon boundaries, and realigned. The consensus sequence of each alignment (defined by a 50% identity threshold up until the putative 3’ end) was extracted, and rare insertions that introduced gaps in the consensus were manually removed. With the 3’ boundary of the IS element, and thus the 3’ boundary of the TnpB ωRNA properly defined, a covariance model of the TnpB ωRNA could be built.

From a randomly selected member of each of the 300 clusters, a 250-bp window of sequence 5’ of the 3’ end of the ωRNA was extracted. A structurally based multiple alignment was then performed using mLocARNA^61^ and used to generate a TnpB-specific ωRNA covariance model with Infernal, refined with CMfinder^60^, and verified with R-scape^63^ at an E-value threshold of 1e-5. This was iterated twice to generate the final generic model of TnpB-associated ωRNA. In addition, more localized ωRNA covariance models were created for each of the 4 TnpB homologs used in this study (*Gst*TnpB1-4). Each protein was used as a seed query in a phmmer (v3.3.2) search against the NR database, with an inclusion and reporting threshold of 1e-30 to identify close relatives of each protein. The steps described above were used to define transposon boundaries and generate ωRNA models using sequences identified in our phmmer search.

#### TnpA detection and autonomous element identification

For both IscB- and TnpB-associated contigs, TnpA was detected using the Pfam Y1_Tnp (PF01797) for a hmmsearch from the HMMR suite (v3.3.2), with an E-value threshold of 1e-4. This search was performed independently on both the curated CDSs of each contig from NCBI and the ORFs predicted by Prodigal^64^ on default settings. The union of these searches was used as the final set of detected TnpA proteins. IS elements that encoded IscB homologs within 1,000 bp of a detected TnpA, or that encoded TnpB homologs within 10,000 bp of a detected TnpA, were defined as autonomous. Analysis which uncovered association with serine resolvases (PF00239) was performed with the same parameters mentioned above.

#### Orientation bias analysis

The closest NCBI-annotated/predicted CDS upstream of each transposon-encoded gene (*tnpB*/*iscB* or the IS*630* transposase) was retrieved and analyzed relative to the gene itself. Initially, the metadata for every NCBI-annotated CDS within contigs containing these genes (*tnpB*/*iscB* or IS*630*) were retrieved, including coordinates and strandedness. Using this information, the closest upstream CDS was identified for each gene based on distance. Then, the annotated orientation of the closest upstream CDS was compared to the annotated orientation of the respective transposon-encoded gene (*tnpB*/*iscB* or IS*630*), to determine whether they were matching. This analysis was performed for gene/CDS pairs at all distances between 0–1000 bp upstream (5’) of the transposon-encoded gene ORF, where 0-bp was defined as overlapping, using a custom Python script.

#### Transposon boundary and TAM/TEM motif determination for *G. stearothermophilus* IS elements (IS*Gst*)

IS*200*/IS*605* elements found in *G. stearothermophilus* strain DSM 458 (NCBI Accession: NZ_CP016552.1) that encoded *iscB* or *tnpB* were identified by a protein homology-based search, as described above. Initial identification of transposon boundaries ware identified by multiple sequence alignment of each unique *tnpB* or *iscB* gene using DNA sequences flanking the TnpB/IscB ORF, and were aligned using MUSCLE^62^ (5.1) PPP algorithm in Geneious (2023.0.1). To build covariance models of the transposon ends, cmfinder was used to detect structural motifs for each end of IS*Gst1*, IS*Gst2* and IS*Gst5* (LE and RE separately) and produce an alignment based on secondary structure. This model was then used for further searches (CMSearch), to identify structurally similar positions within the genome of *G. stearothermophilus* strain DSM 458. All transposon ends were initially paired with the most similar query end and then manually curated, to ensure each the LE and RE within a given pair were correctly positioned relative to each other. This analysis identified several PATE-like elements lacking any protein-coding genes, and a total of 47 IS elements were identified with similar LE and RE sequences. 50 bp upstream and downstream were extracted and aligned using MUSCLE^62^ (5.1) PPP algorithm in Geneious and trimmed using trimAl^56^ (v1.4.rev15), to capture transposon boundaries and identify TAM and TEM motifs based on previous literature describing the location of these essential motifs^41^. Transposon DNA guide regions were predicted based on structural similarities to the transposon ends of *H. pylori* IS*608* and covarying mutations at those predicted locations. TAM motifs, which function as target sites for the transposon insertion event, were confirmed by blastn analysis of DNA sequences flanking predicted transposon boundaries to the NT or WGS database. Phylogenetic trees of transposon ends were built using FastTree^65^ (2.1.11) with default parameters.

#### Small RNA-seq analyses

Small RNA-seq reads were retrieved from NCBI SRA database under accession SRX3260293. Reads were downloaded using the SRA toolkit (2.11.0) and mapped to genomic regions encoding *G. stearothermophilus* IscB and TnpB homologs used in this study, using *G. stearothermophilus* strain ATCC 7953 (GCA_000705495.1) from which small RNA-seq data derives^36^. Reads were mapped using Geneious RNA assembler at medium sensitivity and visualized using Integrative Genomics Viewer^66^.

#### Plasmid construction

All strains and plasmids used in this study are described in **Supplementary Tables 1 and 2**, respectively, and a subset is available from Addgene. In brief, genes encoding TnpA, TnpB, and IscB homologs from *G. stearothermophilus*, *H. pylori* and *D. radiodurans* (**Supplementary Table 3**) were synthesized by GenScript, along mini-Tn elements containing a chloramphenicol resistance gene. To generate mini-Tn plasmids, gene fragments (GenScript) encoding the transposase (TnpA) downstream of a lac and T7 promoter, and transposon ends flanking a chloramphenicol resistance gene, were cloned into EcoRI sites of pUC57. To generate pEffector plasmids, gene fragments (Genscript) of ωRNA encoded downstream of T7 promoter, along with *tnpB* or *iscB* also encoded downstream of T7 promoter, were cloned into pCDF-Duet1 vectors at PfoI and Bsu36I sites. Oligonucleotides containing J23-series promoters were cloned into SalI and KpnI sites, replacing the T7 promoter for ωRNA expression, or into PfoI-XhoI sites, replacing the T7 promoter for *tnpB* expression. pTarget plasmids were generated using a minimal pCOLADuet-1, generated by around-the-horn PCR to create a minimal pCOLA-Duet-1 containing only the ColA origin of replication and kanamycin resistance gene. This vector was then used to generate pTargets encoding 45-bp target sites by around-the-horn PCR. Derivatives of these plasmids were cloned using a combination of methods, including Gibson assembly, restriction digestion-ligation, ligation of hybridized oligonucleotides, and around-the-horn PCR. Plasmids were cloned, propagated in NEB Turbo cells (NEB), purified using Miniprep Kits (Qiagen), and verified by Sanger sequencing (GENEWIZ).

#### Recombineering

Lambda Red (λ-Red) recombination was used to generate genomically integrated mini-Tn cassette. In brief, *E. coli* strain MG1655 (sSL0810) was transformed with pSIM6 (pSL2684) carrying a temperature-sensitive vector encoding λ-Red recombination genes, generating strain sSL2681, and cells were made electrocompetent using standard methods. Fragments for recombineering were generated using standard PCR amplification with primers to generate 50-bp overhangs homologous to the sites of integration. PCR fragments were gel extracted and used to electroporate sSL2681, and cells were recovered for 24 h in LB media. Cells were spun down and plated onto LB-agar containing kanamycin (50 µg ml^−1^) to select for mini-Tn cassette integration. Single colonies were isolated and confirmed to contain a genomically integrated mini-Tn within the *lacZ* locus by colony PCR and Sanger sequencing.

#### Transposon excision assays

For each excision experiment involving a plasmid-based IS element, a single plasmid encoding for TnpA and a chloramphenicol resistance gene-containing mini-Tn IS element was used to transform *E. coli* strain MG1655. Cultures were grown overnight at 37 °C on LB-agar under antibiotic selection (100 µg ml^−1^ carbenicillin, 25 µg ml^−1^ chloramphenicol). Next, three colonies were picked from each agar plate and used to inoculate 5 ml LB supplemented with 0.05 mM IPTG and antibiotic for only for backbone marker (100 µg ml^−1^ carbenicillin). The liquid cultures were incubated at 37 °C for 24 h. Cell lysates were generated, as described previously^18^. In brief, the optical density at 600 nm was measured for liquid cultures. Approximately 3.2 x 10^8^ cells (equivalent to 200 μl of OD_600_ = 2.0) were transferred to a 96-well plate. Cells were pelleted by centrifugation at 4,000g for 5 min and resuspended in 80 μl of H_2_O. Next, cells were lysed by incubating at 95 °C for 10 min in a thermal cycler. The cell debris was pelleted by centrifugation at 4,000g for 5 min, and 10 μl of lysate supernatant was removed and serially diluted with 90 μl of H_2_O to generate 10- and 100-fold lysate dilutions for PCR and qPCR analysis.

IS element excision from the plasmid backbone was detected by PCR using One*Taq* 2X Master Mix with Standard Buffer (NEB) and 0.2 uM primers, designed to anneal upstream and downstream of the IS element. PCR reactions contained 0.5 µl of each primer at 10 µM, 12.5 µl of OneTaq 2X MasterMix with Standard Buffer, 2 µl of 100-fold diluted cell lysate serving as template, and 9.5 µl of H_2_0. The total volume per PCR was 25 µl. Measurements were performed in a BioRad T100 thermal cycler using the following thermal cycling parameters: DNA denaturation (94 °C for 30 s), 35 cycles of amplification (annealing: 52 °C for 20 s, extension: 68 °C for 30 s), followed by a final extension (68 °C for 5 min). Products were resolved by 1.5% agarose gel electrophoresis and visualized by staining with SYBR Safe (Thermo Fisher Scientific). IS element excision events were confirmed by Sanger sequencing of gel-extracted, column-purified (Qiagen) PCR amplicons (GENEWIZ/Azenta Life Sciences).

For excision events involving genomically integrated IS elements, lysate was prepared as described above but harvested from LB-agar containing carbenicillin (100 µg ml^−1^), spectinomycin (100 µg ml^−1^), and X-gal (200 mg ml^−1^) in transposition assays combining TnpA and TnpB, as described below. Measurements were performed in a BioRad T100 thermal cycler using the following thermal cycling parameters: DNA denaturation (94 °C for 30 s), 26 cycles of amplification (annealing: 52 °C for 20 s, extension: 68 °C for 1:15 min), followed by a final extension (68 °C for 5 min). Products were resolved by 1.5% agarose gel electrophoresis and visualized by staining with SYBR Safe (Thermo Fisher Scientific). IS element excision events were confirmed by Sanger sequencing of gel-extracted, column-purified (Qiagen) PCR amplicons (GENEWIZ/Azenta Life Sciences).

#### qPCR quantification of IS element excision

IS element excision frequency from a plasmid backbone was detected by qPCR using SsoAdvanced™ Universal SYBR Green Supermix. qPCR analysis (**Extended Data Figure 3c-e**) was performed using a donor joint-specific primer along with a flanking primer designed to amplify only the excision product; genome-specific primers for relative quantification were designed to amplify the *E. coli* reference gene, *rssA*. 10 µl qPCR reactions containing 5 µl of SsoAdvanced™ Universal SYBR Green Supermix, 2 µl of 2.5 µM primer pair, 1 µl H_2_O, and 2 µl of tenfold-diluted lysate were prepared as described for transposon excision assays. Reactions were prepared in 384-well clear/white PCR plates (BioRad), and measurements were performed on a CFX384 RealTime PCR Detection System (BioRad) using the following thermal cycling parameters to selectively amplify excision products: polymerase activation and DNA denaturation (98 °C for 2.5 min), 40 cycles of amplification (98 °C for 10 s, 62 °C for 20 s), and terminal melt-curve analysis (65–95 °C in 0.5 °C per 5 s increments).

To confirm the sensitivity of qPCR-based measurements from plasmid encoded mini-Tn substrates, we prepared lysates from cells harboring a plasmid containing a mock excised mini-Tn substrate (pSL4826) and a plasmid containing the mini-Tn but lacking an active TnpA transposase required for excision (pSL4735). We simulated variable IS element excision frequencies across five orders of magnitude (ranging from 0.002% to 100%) by mixing cell lysates the control strain and the IS-encoding strain in various ratios, which demonstrated accurate detection of excision products in genomic IS element excision assays *in vivo* to a frequency of 0.001 (**Extended Data Figure 3d**).

Similarly, IS element excision frequencies of genomically integrated mini-TN were quantified by qPCR using SsoAdvanced™ Universal SYBR Green Supermix (BioRad) (**Extended Data Figure 7**). Cells were harvested from LB containing carbenicillin (100 µg ml^−1^), spectinomycin (100 µg ml^−1^), and X-gal (200 mg ml^−1^), as described above. qPCR analysis was performed using transposon flanking- and genome-specific primers. Transposon flanking primers were designed to amplify an approximately 209-bp fragment upon excision. An unexcised product would yield 1,661 bp unexcised fragment. A separate pair of genome-specific primers was designed to amplify an *E. coli* reference gene (*rssA*) for normalization purposes (**Supplementary Table 4**). 10 µl qPCR reactions containing 5 µl of SsoAdvanced™ Universal SYBR Green Supermix, 2 µl of 2.5 µM primer pair, 1 µl H_2_O, and 2 µl of tenfold-diluted lysate were prepared, as described for transposon excision assays. Reactions were prepared in 384-well clear/white PCR plates (BioRad), and measurements were performed on a CFX384 RealTime PCR Detection System (BioRad) using the following thermal cycling parameters to selectively amplify excision products: polymerase activation and DNA denaturation (98 °C for 2.5 min), 40 cycles of amplification (98 °C for 10 s, 60 °C for 20 s), and terminal melt-curve analysis (65–95 °C in 0.5 °C per 5 s increments).

To confirm the sensitivity of qPCR-based measurements from genomically integrated mini-Tn, we prepared lysates from a control MG1655 strain, and a strain containing a genomically-encoded IS element that disrupts the lacZ gene. Similar to the plasmid-based assay, we simulated variable IS element excision frequencies across five orders of magnitude (ranging from 0.002% to 100%) by mixing cell lysates the control strain and the IS-encoding strain in various ratios, and showed that we could accurately detect excision products in genomic IS element excision assays *in vivo* to a frequency of 0.001 (**Extended Data Figure 7b**).

#### Mating-out assays

sSL1592, a kind gift from Dr. Joseph E. Peters, harbors a mini-F plasmid derivative with an integrated spectinomycin cassette. This strain was transformed with a plasmid carrying a mini-Tn harboring a kanamycin marker and either *Gst*TnpA (pSL4245) or catalytically inactive *Gst*TnpA (pSL4974). Cells were selected on LB media containing spectinomycin (100 µg ml^−1^), carbenicillin (100 µg ml^−1^), and kanamycin (50 µg ml^−1^) to generate a donor strain. Three independent colonies were inoculated in liquid LB media containing spectinomycin (100 µg ml^−1^), carbenicillin (100 µg ml^−1^), kanamycin (50 µg ml^−1^), and 0.05 mM IPTG to induce expression of TnpA for 12 h at 37 °C. In parallel, the recipient strain harboring genomically encoded resistance for rifampicin and nalidixic acid were grown in liquid LB media containing rifampicin (100 µg/mL) and nalidixic acid (30 µg/mL) for 12 h at 37 °C. Cells were 100-fold diluted into fresh liquid LB media with respective antibiotics and grown for 2 h to ∼0.5 OD. Cells were then washed with H_2_O and mixed at a concentration of 5 × 107 for both donor and recipient cells, and plated onto solid LB-agar media with no antibiotic selection. Cells were grown for 20 h at 37 °C, scraped off plates, and resuspended in H_2_O. Cells were then serially diluted and plated onto LB media containing rifampicin (100 µg/mL), nalidixic acid (30 µg/mL), spectinomycin (100 µg ml^−1^), and kanamycin (50 µg ml^−1^) to monitor transposition. In addition, cells were also plated to rifampicin (100 µg ml^−1^), nalidixic acid (30 µg ml^−1^), and spectinomycin (100 µg ml^−1^), to determine the entire transconjugant population. The frequency of transposition was calculated by taking the number of colonies that exhibited Nal^R^ + Rif^R^ + Spec^R^ + Kan^R^ phenotype (i.e. transposition positive), divided by the number of transconjugants that exhibited a Nal^R^ + Rif^R^ + Spec^R^ phenotype. Transconjugants showing resistance to nalidixic acid, rifampicin, spectinomycin, and kanamycin were isolated using Zymo Research ZR BAC DNA miniprep kit and sequenced using nanopore long-read sequencing (Plasmidsaurus). Reads were analyzed in Geneious Prime (2023.0.1) by using a custom blast database to identify reads containing mini-Tn and flanking mini-F plasmid sequence. Insertion events were aligned to Mini-F plasmid reference to identify sites of integration.

#### Plasmid interference assays

Plasmid interference assays were performed in *E. coli* BL21 (DE3) (**Fig. 3c,f and Extended Data Figure 5a,b,d**) or *E coli* str. K-12 substr. MG1655 (sSL0810) strains for all other experiments. For **Figure 3c** (TnpB homologs), BL21 (DE3) cells were transformed with pTarget plasmids, and single colony isolates were selected to prepare chemically competent cells. 400 ng of pEffector plasmids were then delivered via transformation. After 3 h, cells were spun down at 4000 *g* for 5 min and resuspended in 20 µl of H_2_O. Cells were then serial diluted (10x) and transferred to LB media containing spectinomycin (100 µg ml^−1^), kanamycin (50 µg ml^−1^), and 0.05 mM IPTG and grown for 24 h at 37 °C. For all remaining spot assays using MG1655 strains, chemically competent cells were first prepared with pEffector plasmid and then transformed with 400 ng of pTarget plasmids. After 3 h, cells were spun down at 4000 *g* for 5 min and resuspended in 20 µl of H_2_O. Cells were then serial diluted (10x) and transferred to LB media containing spectinomycin (100 µg ml^−1^), kanamycin (50 µg ml^−1^), and 0.05 mM IPTG and grown for 14 h at 37 °C. Plates were imaged in an Amersham Imager 600.

Quantification of plasmid interference was calculated by determining the number of colony forming units (CFUs) following transformation. Cells were first transformed with pEffector plasmids and prepped as chemically competent cells for a second round of transformation with 200 ng of pTarget. Cells were then spun down at 4000 *g* for 5 min and resuspended in 100 μL of H_2_O. Cells were then serial diluted and plated to LB media containing spectinomycin (100 µg ml^−1^), kanamycin (50 µg ml^−1^). 0.05 mM IPTG was added to media when T7 promoter was used. CFUs were counted following 24 h of growth at 37 °C. Frequencies were normalized relative to a non-targeting guide RNA.

#### Genome targeting and cell killing assays

Cell killing assays via genomic targeting with TnpB (**Figure 3h and Extended Data Figure 5e**) or IscB (**Figure 3h**) were performed by transforming *E. coli* str. K-12 substr. MG1655 (sSL0810) strains with spectinomycin-resistant plasmids constitutively expressing TnpB/IscB and either genomic targeting or non-targeting guide RNAs. Cells were transformed with 400 ng plasmid. After 3 h, cells were spun down at 4000 *g* for 5 min and resuspended in 20 µl of H_2_O. Cells were then serial diluted (10x) and transferred to LB media containing spectinomycin (100 µg ml^−1^) and grown for 24 h at 37 °C.

#### ChIP-seq experiments and library preparation

ChIP-seq experiments were generally performed as described previously^67^. The following active site mutations were introduced to inactivate the endonuclease domains of the respective 3×Flag-tagged proteins to simulate DNA binding prior to DNA cleavage: GstIscB (D87A, H238A, H239A); GstTnpB (D196A); SpyCas9 (D10A, H840A); AsCas12a (D908A). *E. coli* BL21(DE3) cells were transformed with a single plasmid encoding the catalytically inactive effector and either a *lacZ* targeting ωRNA or non-targeting ωRNA. After incubation for 16 h at 37°C on LB agar plates with antibiotics (200 µg ml^−1^ spectinomycin), cells were scraped and resuspended in 1 ml of LB. The optical density at 600 nm (OD600) was measured, and approximately 4.0 × 10^8^ cells (equivalent to 1 ml with an OD600 of 0.25) were spread onto two LB agar plates containing antibiotics (200 µg ml^−1^ spectinomycin) and supplemented with 0.05 mM IPTG. Plates were incubated at 37°C for 24 h. All cell material from both plates was scraped and transferred to a 50 ml conical tube.

Cross-linking was performed by mixing 1 ml of formaldehyde (37% solution; Thermo Fisher Scientific) to 40 ml of LB medium (∼1% final concentration) followed by immediate resuspension of the scraped cells by vortexing and 20 min of gentle shaking at room temperature. Cross-linking was stopped by the addition of 4.6 ml of 2.5 M glycine (∼0.25 M final concentration) followed by 10 min incubation with gentle shaking. Cells were pelleted at 4°C by centrifuging at 4,000g for 8 min. The following steps were performed on ice using buffers that had been sterile-filtered. The supernatant was discarded, and the pellets were fully resuspended in 40 ml TBS buffer (20 mM Tris-HCl pH 7.5, 0.15 M NaCl). After centrifuging at 4,000 *g* for 8 min at 4 °C, the supernatant was removed, and the pellet was resuspended in 40 ml TBS buffer again. Next, the OD600 was measured for a 1:1 mixture of the cell suspension and fresh TBS buffer, and a standardized volume equivalent to 40 ml of OD600 = 0.6 was aliquoted into new 50 ml conical tubes. A final 8 min centrifugation step at 4,000 *g* and 4 °C was performed, cells were pelleted and the supernatant was discarded. Residual liquid was removed, and cell pellets were flash-frozen using liquid nitrogen and stored at −80 °C or kept on ice for the subsequent steps.

Bovine serum albumin (GoldBio) was dissolved in 1× PBS buffer (Gibco) and sterile-filtered to generate a 5 mg ml−1 BSA solution. For each sample, 25 μl of Dynabeads Protein G (Thermo Fisher Scientific) slurry (hereafter, beads or magnetic beads) were prepared for immunoprecipitation. Up to 250 μl of the initial bead slurry were prepared in a single tube, and washes were performed at room temperature, as follows: the slurry was transferred to a 1.5 ml tube and placed onto a magnetic rack. The supernatant was removed, 1 ml BSA solution was added, and the beads were fully resuspended by vortexing, followed by rotating for 30 s. This was repeated for three more washes. Finally, the beads were resuspended in 25 μl (× n samples) of BSA solution, followed by addition of 4 μl (× n samples) of monoclonal anti-Flag M2 antibodies produced in mouse (Sigma-Aldrich). The suspension was moved to 4 °C and rotated for >3 h to conjugate antibodies to magnetic beads. While conjugation was proceeding, cross-linked cell pellets were thawed on ice, resuspended in FA lysis buffer 150 (50 mM HEPES-KOH pH 7.5, 0.1% (w/v) sodium deoxycholate, 0.1% (w/v) SDS, 1 mM EDTA, 1% (v/v) Triton X-100, 150 mM NaCl) with protease inhibitor cocktail (Sigma-Aldrich) and transferred to a 1 ml milliTUBE AFA Fiber (Covaris). The samples were sonicated on a M220 Focused-ultrasonicator (Covaris) with the following SonoLab 7.2 settings: minimum temperature, 4 °C; set point, 6 °C; maximum temperature, 8 °C; peak power, 75.0; duty factor, 10; cycles/bursts, 200; 17.5 min sonication time. After sonication, samples were cleared of cell debris by centrifugation at 20,000 *g* and 4 °C for 20 min. The pellet was discarded, and the supernatant (∼1 ml) was transferred into a fresh tube and kept on ice for immunoprecipitation. For non-immunoprecipitated input control samples, 10 μl (∼1%) of the sheared cleared lysate were transferred into a separate 1.5 ml tube, flash-frozen in liquid nitrogen and stored at −80 °C.

After >3 h, the conjugation mixture of magnetic beads and antibodies was washed four times with BSA solution as described above, but at 4 °C. Next, the beads were resuspended in 30 μl (× n samples) FA lysis buffer 150 with protease inhibitor, and 31 μl of resuspended antibody-conjugated beads were mixed with each sample of sheared cell lysate. The samples rotated overnight for 12–16 h at 4 °C for immunoprecipitation of Flag-tagged proteins. The next day, tubes containing beads were placed on a magnetic rack, and the supernatant was discarded. Then, six bead washes were performed at room temperature, as follows, using 1 ml of each buffer followed by sample rotation for 1.5 min: (1) two washes with FA lysis buffer 150 (without protease inhibitor); (2) one wash with FA lysis buffer 500 (50 mM HEPES-KOH pH 7.5, 0.1% (w/v) sodium deoxycholate, 0.1% (w/v) SDS, 1 mM EDTA, 1% (v/v) Triton X-100, 500 mM NaCl); (3) one wash with ChIP wash buffer (10 mM Tris-HCl pH 8.0, 250 mM LiCl, 0.5% (w/v) sodium deoxycholate, 0.1% (w/v) SDS, 1 mM EDTA, 1% (v/v) Triton X-100, 500 mM NaCl); and (4) two washes with TE buffer 10/1 (10 mM Tris-HCl pH 8.0, 1 mM EDTA). The beads were then placed onto a magnetic rack, the supernatant was removed, and the beads were resuspended in 200 μl of fresh ChIP elution buffer (1% (w/v) SDS, 0.1 M NaHCO3). To release protein–DNA complexes from beads, the suspensions were incubated at 65 °C for 1.25 h with gentle vortexing every 15 min to resuspend settled beads. During this incubation, the non-immunoprecipitated input samples were thawed, and 190 μl of ChIP Elution Buffer was added, followed by the addition of 10 μl of 5 M NaCl. After the 1.25 h incubation of the immunoprecipitated samples was complete, the tubes were placed back onto a magnetic rack, and the supernatant containing eluted protein–DNA complexes was transferred to a new tube. Then, 9.75 μl of 5 M NaCl was added to ∼195 μl of eluate, and the samples (both immunoprecipitated and non-immunoprecipitated controls) were incubated at 65 °C overnight to reverse-cross-link proteins and DNA. The next day, samples were mixed with 1 μl of 10 mg ml^−1^ RNase A (Thermo Fisher Scientific) and incubated for 1 h at 37 °C, followed by addition of 2.8 μl of 20 mg ml^−1^ proteinase K (Thermo Fisher Scientific) and 1 h incubation at 55 °C. After adding 1 ml of buffer PB (QIAGEN recipe), the samples were purified using QIAquick spin columns (QIAGEN) and eluted in 40 μl TE buffer 10/0.1 (10 mM Tris-HCl pH 8.0, 0.1 mM EDTA).

ChIP–seq Illumina libraries were generated for immunoprecipitated and input samples using the NEBNext Ultra II DNA Library Prep Kit for Illumina (NEB). Sample concentrations were determined using the DeNovix dsDNA Ultra High Sensitivity Kit. Starting DNA amounts were standardized such that an approximately equal mass of all input and immunoprecipitated DNA was used for library preparation. After adapter ligation, PCR amplification (12 cycles) was performed to add Illumina barcodes, and ∼450 bp DNA fragments were selected using two-sided AMPure XP bead (Beckman Coulter) size selection, as follows: the volume of barcoded immunoprecipitated and input DNA was brought up to 50 μl with TE Buffer 10/0.1; in the first size-selection step, 0.55× AMPure beads (27.5 μl) were added to the DNA, the sample was placed onto a magnetic rack, and the supernatant was discarded and the AMPure beads were retained; in the second size-selection step, 0.35× AMPure beads (17.5 μl) were added to the DNA, the sample was placed onto a magnetic rack, and the AMPure beads were discarded and the supernatant was retained. The concentration of DNA was determined for pooling using the DeNovix dsDNA High Sensitivity Kit.

Illumina libraries were sequenced in paired-end mode on the Illumina MiniSeq and NextSeq platforms with automated demultiplexing and adapter trimming (Illumina). For each ChIP–seq sample, >1,000,000 raw reads (including genomic and plasmid-mapping reads) were obtained.

#### ChIP-seq data analyses

ChIP-seq data analysis was generally performed as described previously^67^. In brief, ChIP-seq paired-end reads were trimmed and mapped to an *E. coli* BL21(DE3) reference genome (GenBank: CP001509.3). Genomic *lacZ* and *lacI* regions, partially identical to plasmid-encoded genes, were masked in all alignments (genomic coordinates: 335,600–337,101 and 748,601–750,390). In the ChIP-seq analysis of Cas9 and Cas12a, rrnB t1 terminator genomic sequence was masked (genomic coordinates: 4,121,275-4,121,400). Mapped reads were sorted, indexed, and multi-mapping reads were excluded. Aligned reads were normalized by RPKM and visualized in IGV^66^. For genome-wide views, maximum read coverage values were plotted in 1-kb bins. Peak calling was performed using MACS3^68^ with respect to non-immunoprecipitated control samples of TnpB and Cas9. The peak summit coordinates in the MACS3 output summits.bed file were extended to encompass a 200-bp window using BEDTools^69^. The corresponding 200-bp sequence for each peak was extracted from the E. coli reference genome using the command bedtools getfasta. Sequence motifs were determined using MEME ChIP^70^. Individual off-target sequences (**Extended Data Figure 6**) represent sequences from the top enriched peaks determined by MACS3 that contain the MEME ChIP motif.

#### TAM library cloning

TAM libraries were cloned containing a 6-bp randomized sequence between the native target sequences for *Gst*IscB (IS*Gst5*) and *Gst*TnpB2 (IS*Gst2*). In brief, two partially overlapping oligos (oSL9404 and oSL9405) were annealed by heating to 95 °C for 2 min and then cooled to room temperature. One of these oligos (oSL9404) contained a 6-nt degenerate sequence flanked by target sites for *Gst*TnpB2 and *Gst*IscB. Annealed DNA was treated with DNA Polymerase I, Large (Klenow) Fragment (NEB) in 40 µL reactions and incubated at 37 °C for 30 min, then gel purified (QIAGEN Gel Extraction Kit). Double-stranded insert DNA and vector backbone (pSL4031) was digested with BamHI and HindIII (37 °C, 1 h). The digested insert was cleaned-up (Qiagen MinElute PCR Purification Kit), and digested backbone was gel-purified (Qiagen QIAquick Gel Extraction Kit). The backbone and insert were ligated with T4 DNA Ligase (NEB). Ligation reactions were transformed in with electrocompetent NEB 10-beta cells according to the manufacturer’s protocol. After recovery (37 °C for 1 h), cells were plated on large bioassay plates containing LB agar and kanamycin (50 µg ml^−1^). Approximately 5 million CFUs were scraped from each plate, representing 1000x coverage of each library member, and plasmid DNA was isolated using the Qiagen CompactPrep Midi Kit.

#### TAM library assays and NGS library prep

DNA solutions containing 500 ng of the TAM plasmid library (pSL4841) and 500 ng of plasmids encoding either *Gst*TnpB2 (pSL4369) or *Gst*IscB (pSL4514) were co-transformed in electrocompetent *E. coli* BL21(DE3) cells according to the manufacturer’s protocol (Sigma-Aldrich). Cells were serially diluted on large bioassay plates containing LB agar, spectinomycin (100 µg ml^−1^), and kanamycin (50 µg ml^−1^). Approximately 600,000 CFUs were scraped from plates, representing 100x coverage of each library member, and plasmid DNA was isolated using the Qiagen CompactPrep Midi Kit. Illumina amplicon library for NGS was prepared through 2-step PCR amplification. In brief, ∼50 ng of plasmid DNA recovered from TAM assay was used in each “PCR-1” amplification reaction with primers flanking the degenerate TAM library sequence and containing universal Illumina adaptors as 5’ overhangs. Amplification was carried out using high-Fidelity Q5 DNA Polymerase (NEB) for 16 thermal cycles. Samples from “PCR-1” amplification were diluted 20-fold and amplified for “PCR-2” in 10 thermal cycles with primers contain indexed p5/p7 sequences. Reactions were verified by analytical gel electrophoresis. Sequencing was performed with a paired-end run using a MiniSeq High Output Kit with 150-cycles (Illumina).

#### Analyses of NGS TAM library data

Analysis of TAM depletion library was performed using a custom Python script. Demultiplexed reads were filtered to remove reads that did not contain a perfect match to the 58-bp sequence upstream of the degenerate sequence for any i5-reads. For reads that passed this filtering step, the 6-nt degenerate sequence was extracted and counted. The relative abundance of each degenerate sequence in a sample was determined by dividing the degenerate sequence count by the total number of sequence counts for that sample. Then, the fold-change between the output and input libraries was calculated by dividing the relative abundance of each degenerate sequence in the output library by its relative abundance in the input library, and then log2-transformed. Sequence logos were constructed by taking the 10 most depleted sequences and generated using WebLogo^71^ (v2.8).

#### Transposition assays combining TnpA and TnpB

*E. coli* str. K-12 substr. MG1655 (sSL0810) was engineered to carry a genomic integrated mini-Tn containing a kanamycin resistance cassette inserted into *lacZ* by recombineering as described above to generate sSL2771. This strain was transformed with either pCDFDuet-1 (pSL0007) or various *Gst*TnpB carrying vectors (pSL4369, pSL4664, pSL4518 and pSL4740, see **Supplementary Table 2** for description) and selected on LB agar containing spectinomycin (100 µg ml^−1^) and kanamycin (50 µg ml^−1^). Single colony isolates of cells harboring each plasmid were prepared chemically competent and transformed with a TnpA expression vector (pSL4529) or a catalytically inactive mutant TnpA expression vector (pSL4534) and selected on LB agar containing carbenicillin (100 µg ml^−1^), spectinomycin (100 µg ml^−1^) and kanamycin (50 µg ml^−1^). Three single colony isolate of each transformant were grown in liquid LB containing carbenicillin (100 µg ml^−1^), spectinomycin (100 µg ml^−1^) and kanamycin (50 µg ml^−1^) and grown for 14 h at 37 °C. Optical density (OD) of each culture was measured and approximately 10^7^ cells were plated onto MacConkey agar media containing carbenicillin (100 µg ml^−1^), spectinomycin (100 µg ml^−1^) and 0.05mM IPTG for TnpA induction. Importantly, the media did not contain kanamycin to allow for excision of the mini-Tn. Cells were grown at 37 °C for 4 days on MacConkey media to enrich for mini-Tn excision events. Cells were then harvested, serially diluted, and plated onto LB agar containing carbenicillin (100 µg ml^−1^), spectinomycin (100 µg ml^−1^) and X-gal (200 mg ml^−1^) or carbenicillin (100 µg ml^−1^), spectinomycin (100 µg ml^−1^), kanamycin (50 µg ml^−1^) and X-gal (200 mg ml^−1^) and grown for 18 h at 37 °C. Total number of colonies were counted, along with the number of blue colonies to determine the frequency of excision and reintegration events. In addition, genomic lysate was harvested from cells as described above for PCR analysis.

#### Statistics and reproducibility

qPCR and analytical PCRs resolved by agarose gel electrophoresis gave similar results in three independent replicates. Sanger sequencing of excision products was performed once for each isolate. Next-generation sequencing of PCR amplicons was performed once. Plasmid interference assays were performed in three independent replicates. Transposition assays combining TnpA and TnpB were performed with three independent replicates.

#### Data availability

Next-generation sequencing data are available in the National Center for Biotechnology Information (NCBI) Sequence Read Archive: SRX19058888-SRX19058905, SRR23476356-SRR23476358 (BioProject Accession: PRJNA925099) and the Gene Expression Omnibus (GSE223127). The published genome used for ChIP-seq analyses was obtained from NCBI (GenBank: CP001509.3). The published genome used for bioinformatics analyses of the *Geobacillus stearothermophilus* genome was obtained from NCBI (GenBank: NZ_CP016552.1). Datasets generated and analyzed in the current study are available from the corresponding author upon reasonable request.

#### Code availability

Custom scripts used for bioinformatics, TAM library analyses, and ChIP-seq data analyses are available at GitHub (https://github.com/sternberglab/Meers_et_al_2023).

## SUPPLEMENTARY INFORMATION

Full alignments of all *Gst* IS*200/*IS*605* left and right ends used in this study are provided in Supplementary Figure 1 and 2. Supplementary Tables 1 and 2 provides a list of all strains and plasmids used in this study, respectfully. A list of genes and associated NCBI accession numbers used in this study is provided in Supplementary Table 3. Primer pairs used for PCR or qPCR-based assays of IS excision are provided in Supplementary Table 4. ChIP-seq metadata information is provided in Supplementary Table 5.

## ACKNOWLEDGMENTS

We thank D.R. Gelsinger and A. Bernheim for helpful bioinformatics discussions, L.E. Berchowitz for helpful RNA biology discussions, J.C. Cheong for technical support, L.F. Landweber for qPCR instrument access, and the JP Sulzberger Columbia Genome Center for NGS support. C.M. was supported by NIH Postdoctoral Fellowship F32 GM143924-01A1. M.W.G.W was supported by the National Science Foundation (GRFP to M.W.G.W.). This research was supported by a generous start-up package from the Columbia University Irving Medical Center Dean’s Office and the Vagelos Precision Medicine Fund (S.H.S.).

## AUTHOR CONTRIBUTIONS

C.M. and S.H.S. conceived of and designed the project. C.M. performed most experiments, with assistance from S.P. on plasmid interference assays, and F.T.H. for transposon excision assays and ChIP-seq experiments and analyses. C.M. and H.L. performed all bioinformatics analyses. M.W.G.W. assisted with TAM library design and NGS analysis. J.G. assisted with plasmid interference assays. C.M. and S.H.S. discussed the data and wrote the manuscript, with input from all authors.

## SUPPLEMENTARY FIGURES

**Extended Data Figure 1.**
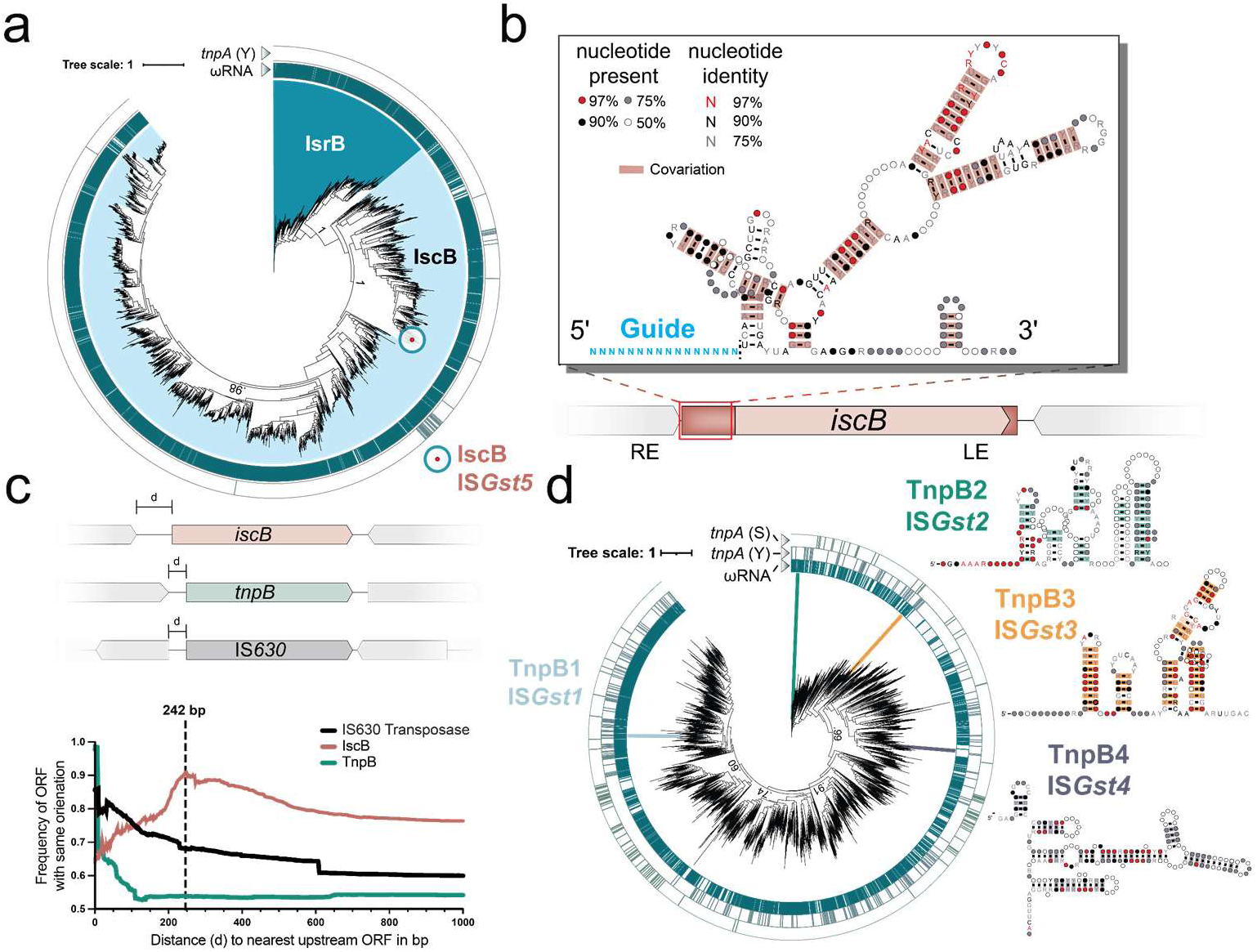
Bioinformatic analyses of IscB and TnpB homologs. **a,** Phylogenetic tree of IscB and IsrB protein homologs; IscB contain HNH and RuvC nuclease domains, whereas IsrB lacks the HNH nuclease. Genetic neighborhood analyses demonstrate that most homologs are encoded proximal to a predicted ωRNA (inner ring), whereas the vast majority do not reside near a predicted TnpA transposase gene (outer ring). The *Gst*IscB homolog used in this study is indicated. Bootstrap values are indicated for major nodes. **b,** Schematic of a non-autonomous IS element encoding IscB and its associated ωRNA; a structural covariation model is shown in the inset. The red rectangle and dotted black line indicate the transposon boundaries, and the guide portion of the ωRNA is shown in blue. LE and RE, transposon left end and right end. **c,** Orientation bias of the nearest upstream ORFs to the indicated protein-coding gene (*iscB*, *tnpB* or IS*630*), demonstrating that IS elements encoding IscB are preferentially integrated (or retained) in an orientation matching that of the upstream gene. The y-axis indicates the frequency of ORFs containing the same orientation, at a distance from the gene start codon defined by the x-axis. 242 bp represents the average length of IscB-associated ωRNAs upstream of IscB ORF. The spike at ∼0-bp for TnpB corresponds to IS elements that encode adjacent/overlapping *tnpA* and *tnpB* genes. IS*630* transposase genes are included as a representative gene from unrelated transposable elements. **d,** Phylogenetic tree of TnpB homologs. Genetic neighborhood analyses demonstrate that most homologs are encoded proximal to a predicted ωRNA (inner ring), whereas the vast majority do not reside near a predicted TnpA transposase gene (outer rings). Bootstrap values are indicated for major nodes. Interestingly, TnpB homologs are associated with two unrelated transposase families, tyrosine transposases (TnpA (Y)) and and serine transposases (TnpA (S)) in bacteria. *Gst*TnpB homologs used in this study are highlighted, along with the predicted structures of their associated ωRNAs, based on covariance modeling. IS*Gst1* TnpB1 was not experimentally active and ωRNA did not show strong covariation in structure and was therefore omitted.

**Extended Data Figure 2.**
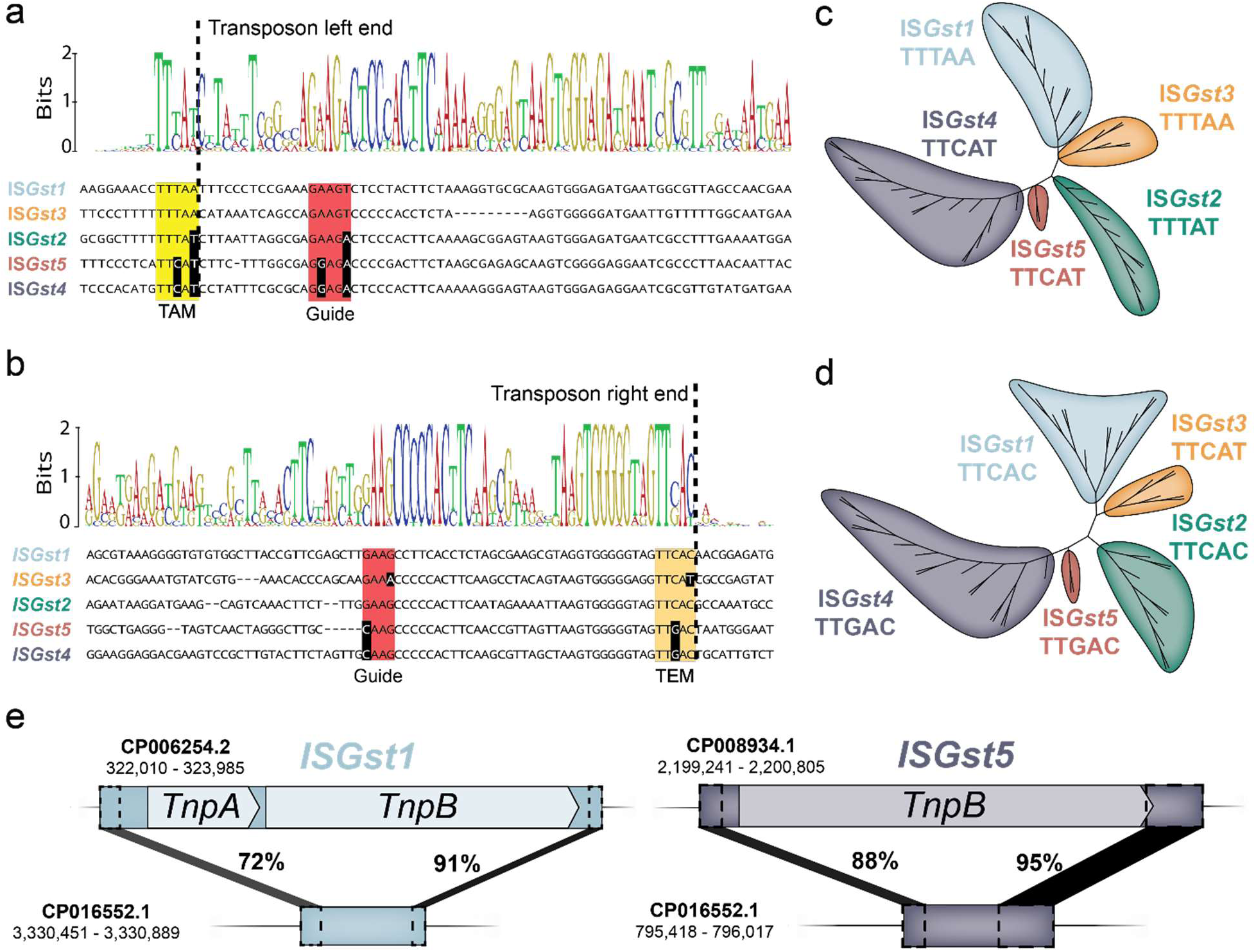
Classification of IS*605*-family elements encoded by *G. stearothermophilus* strain DSM458. **a,** DNA multiple sequence alignment of transposon left ends for IS*200*/IS*605*-family elements from *G. stearothermophilus*. The weblogo (top) is built from 47 unique elements (Supplementary Fig. 1 & 2), and one representative sequence from each family is shown below, with the TAM shown in yellow and DNA guide sequences shown in red as indicated. Nucleotides highlighted in black exhibit covarying mutations, relative to IS*Gst1*. TAM, transposon-adjacent motif; dotted black line indicates the transposon boundary. **b,** DNA multiple sequence alignment of transposon right ends for IS*200*/IS*605*-family elements from *G. stearothermophilus*, shown as in **a**. TEM, transposon-encoded motif is shown in orange. **c,** Phylogenetic tree of IS*Gst* elements based on the transposon left end. Each colored clade encodes an associated TnpB/IscB protein homolog and is flanked by the indicated TAMs sequence. **d,** Phylogenetic tree of IS*Gst* elements based on the transposon right end, shown as in **b** but with TEM sequence in lieu of TAM. **e,** Schematic of PATEs (palindrome associated transposable elements) related to IS*Gst1* and IS*Gst5*, which contain similar transposon ends but no protein-coding genes. The percent sequence identity between shaded regions (black) is shown, as are the genomic accession IDs and coordinates.

**Extended Data Figure 3.**
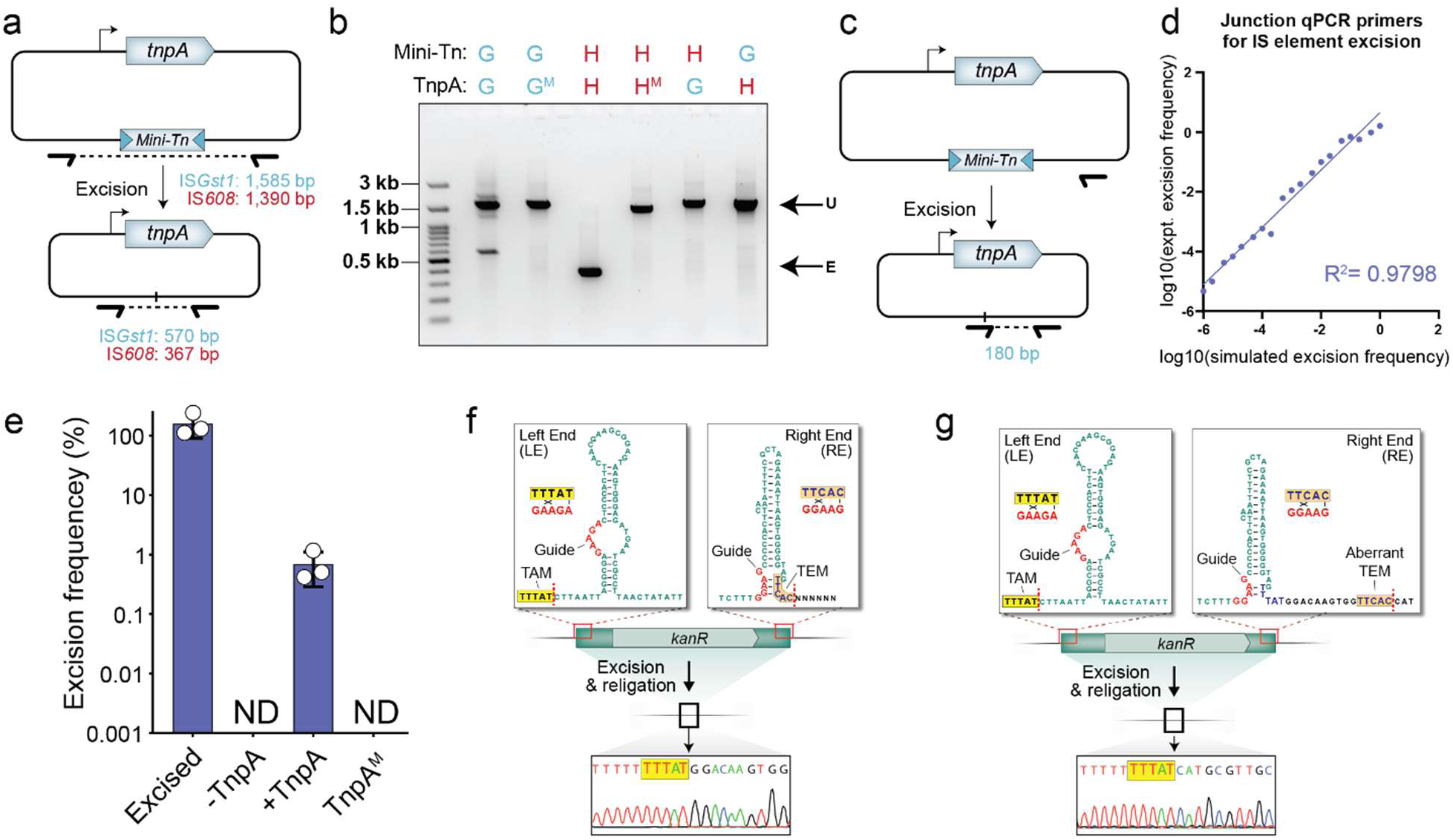
Specificity and efficiency of transposon DNA excision by TnpA. **a,** Schematic of heterologous transposon excision assay in *E. coli*. Plasmids encode TnpA and mini-transposon (Mini-Tn) substrates, whose loss is monitored by PCR using the indicated primers. The expected sizes of PCR products generated from donor joints that are produced upon religation of flanking sequences are shown, for both IS*Gst1* and *H. pylori* IS*608*. **b,** TnpA homologs do not cross-react with distinct IS elements, as assessed by analytical PCR. Cell lysates were tested after overnight expression of TnpA in combination with a mini-Tn substrate, from either *G. stearothermophilus* (G) or *H. pylori* (H), and PCR products were resolved by agarose gel electrophoresis. M refers to catalytically inactive mutants. Note that *Hpy*TnpA is substantially more active for DNA excision than *Gst*TnpA under the tested conditions. U, unexcised; E, excised. **c,** Schematic of qPCR assay to quantify excision frequencies, in which one of the two primers anneals directly to the donor joint formed upon mini-Tn excision and religation. **d,** Comparison of simulated excision frequencies, generated by mixing clonally excised and unexcised lysate in known ratios, versus experimentally determined integration efficiencies measured by qPCR. **e,** qPCR-based quantification of TnpA-mediated excision of an IS*Gst1* mini-Tn substrate in *E. coli*. Mock refers to a cloned excision product; M denotes a TnpA mutant (Y125A); ND, not detected above a 0.0001% threshold. Bars indicate mean ± standard deviation (n = 3). **f,** Schematic of mini-Tn IS*Gst2* element, highlighting the subterminal palindromic transposon ends located on the top strand (top). Transposon-adjacent and transposon-encoded motifs (TAM and TEM) are shown in yellow and orange, respectively; DNA guides are shown in red, and their putative base-pairing interactions are indicated; dotted lines indicate transposon boundaries and thus the sites of ssDNA cleavage and religation. Sanger sequencing of excision events confirm the identity of the expected donor joint product formed upon transposon loss (bottom). Sanger sequencing results are duplicated from Fig. 2D**. g,** Schematic and Sanger sequencing data as in **f**, but for a modified IS*Gst2* substrate containing TEM mutations. Experimentally detected products erroneously excise at an alternative TEM-like sequence located outside of the native transposon boundary (orange), presumably because of the need to maintain cognate base-pairing between the DNA guide and TEM.

**Extended Data Figure 4.**
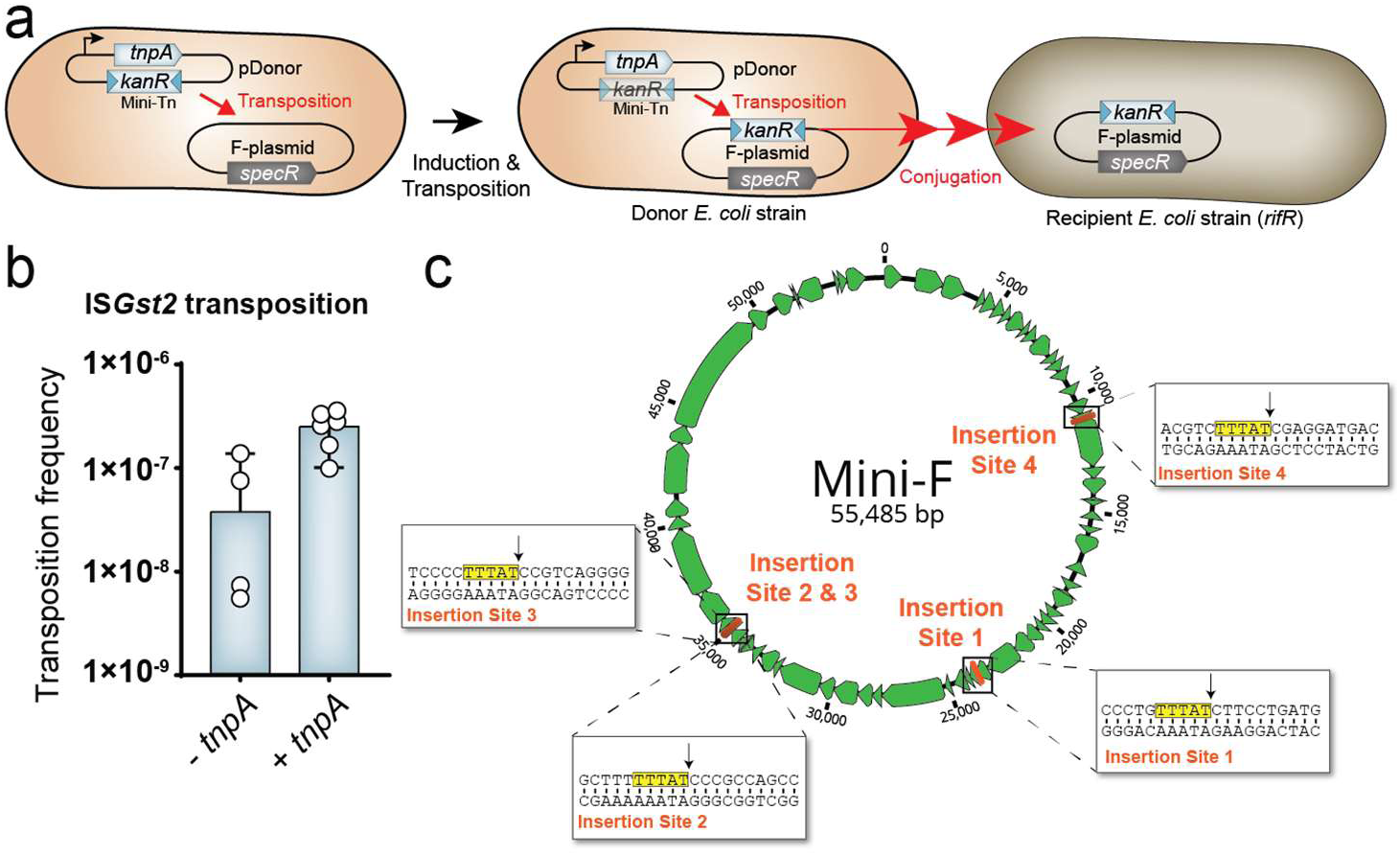
Mating-out assay to monitor transposition of IS*Gst2*. **a,** Schematic of mating-out assay, in which transposition events into the F-plasmid are monitored via drug selection. *E. coli* donor cells carrying an F-plasmid were transformed with a plasmid encoding TnpA and IS*Gst2*-derived mini-Tn. After induction of TnpA, conjugation was used to transfer the F-plasmid into the recipient strain, and transposition events were quantified by selecting for recipient cells (Rif^R^) containing spectinomycin (F^+^) and kanamycin (mini-Tn^+^) resistance. **B,** Transposition frequency of IS*Gst2* into the F-plasmid was measured with and without *tnpA*. Bars indicate mean ± standard deviation (n = 6). **C,** Drug-selected cells from mating-out assays contain TAM-proximal IS insertions, as evidenced by long-read Nanopore sequencing. A genetic map of the F-plasmid is shown, along with the location of distinct IS*Gst2*-derived mini-Tn integration events. The insets show a zoom-in view of each integration site at the nucleotide level, with the TAM motif highlighted in yellow and the integration site specified by an arrow.

**Extended Data Figure 5.**
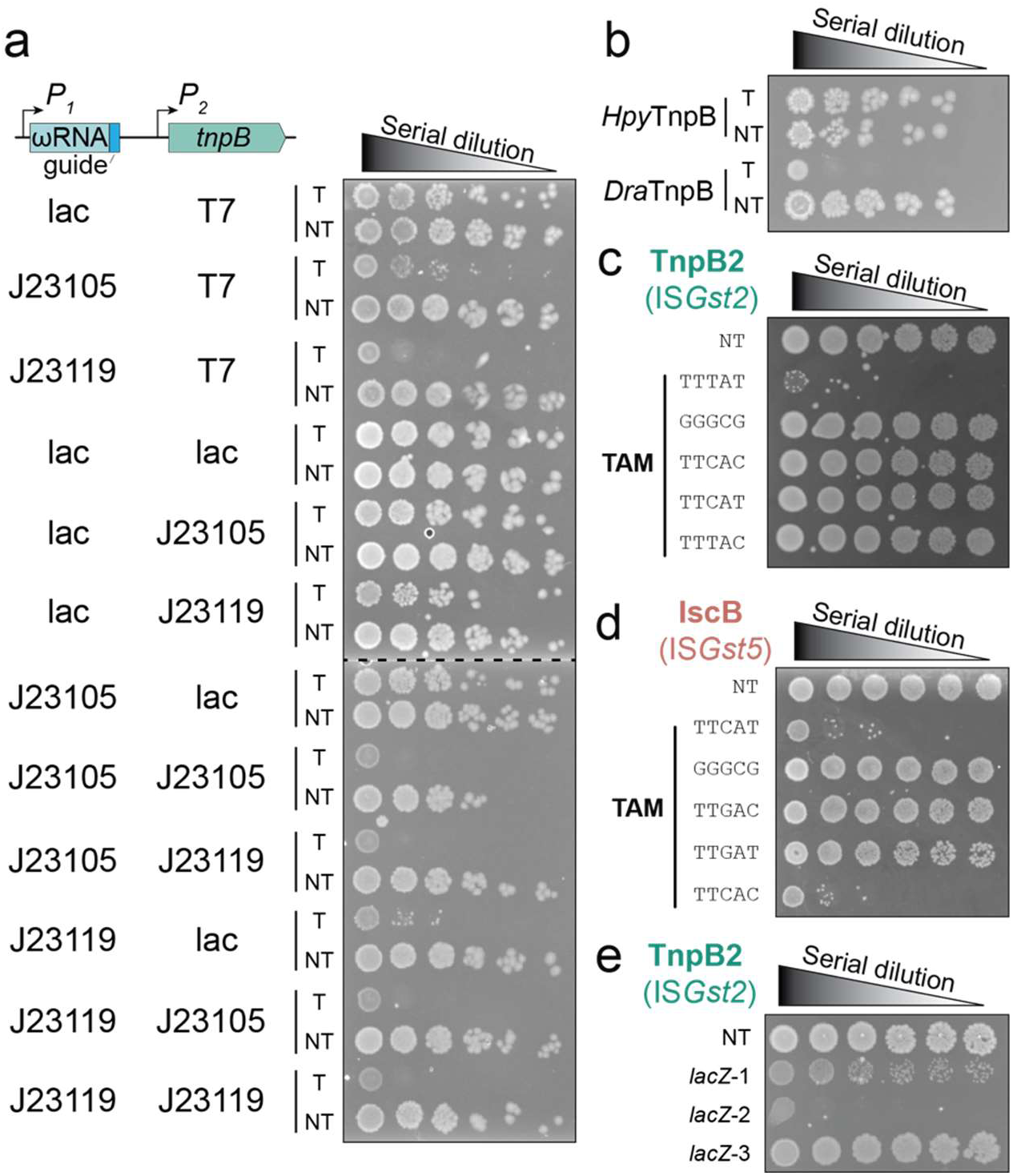
Optimization and testing of DNA cleavage parameters with TnpB/IscB nucleases. **a,** Promoter screen to optimize conditions for *E. coli*-based interference assays using plasmid-encoded ωRNA and TnpB2. P1 indicates promoters for ωRNA expression, P2 indicates promoters for TnpB2 expression. Transformants with a targeting (T) or non-targeting (NT) ωRNA-pTarget combination were serially diluted and plated on selective media at 37 °C for 24 h. **b,** Results from plasmid interference assays with *Hpy*TnpB (IS*608*) and *Dra*TnpB (IS*Dra2*) using ωRNAs that target native donor joint products, which revealed an absence of activity for *Hpy*TnpB. Experiments were performed as in **a**. **c,** DNA cleavage by TnpB2 is highly sensitive to TAM mutations, as assessed by plasmid interference assays. Data are shown as in **a**, with the indicated TAM sequences; TTTAT denotes the WT TAM, and NT denotes a non-targeting control. **d,** DNA cleavage by IscB is highly sensitive to TAM mutations, as assessed by plasmid interference assays. Data are shown as in **a**, with the indicated TAM sequences; TTCAT denotes the WT TAM, and NT denotes a non-targeting control. **e,** TnpB2 is only active for targeted genomic DNA cleavage using select ωRNAs, as assessed by genome targeting assays. Transformants with a non-targeting (NT) or one of three *lacZ*-specific guides were serially diluted and plated on selective media at 37 °C for 24 h.

**Extended Data Figure 6.**
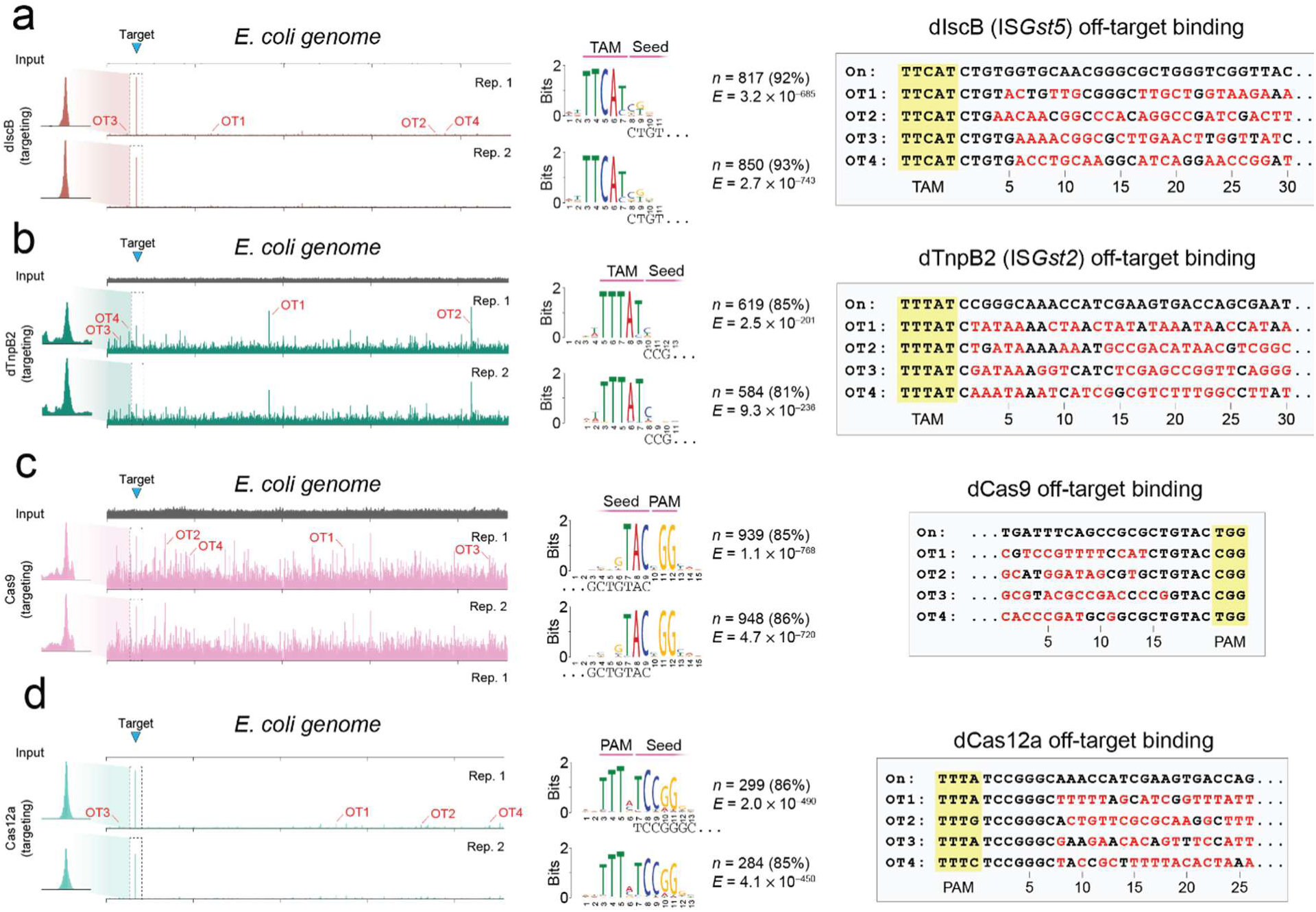
Off-target ChIP-seq DNA binding analyses. **a,** ChIP-seq experiments reveal recruitment of dIscB to the target site (blue triangle) with a targeting ωRNA shown as two independent reps. Genome-wide representation of ChIP-seq data for dIscB reshown from figure 4B with addition of second replicate. Representative off-target sites for dIscB identified by MACS3 are highlighted (OT1-4) and analyzed in middle and right panels, respectively. Middle panel highlights analysis of off-target binding events by dIscB using MEME ChIP, as shown in Fig. 4B. Motifs shared by off-target peaks reveal conserved TAM sequences and little conservation of the adjacent seed sequence (left). The sequence of the 5’ end of the corresponding ωRNA is shown at the bottom of each motif. Two targeting replicates are shown. n indicates the number of peaks contributing to the motif and their percentage of total peaks called by MACS3; E, E-value significance of the motif generated from the MEME ChIP analysis (right of weblogo). DNA sequences corresponding to the on-target and off-target sites are shown on right with TAM (yellow) and mismatches (red) highlighted. OT1-4 represent the top enrichment peaks contributing to each motif, as called by MACS3 with respect to the input sample (Methods). **b,** ChIP-seq experiments reveal recruitment of dTnpB2 to the target site (blue triangle) with a targeting ωRNA shown as two independent replicates. Data shown as in **a**. Similar to dIscB, dTnpB2 shows limited seed sequence requirements. **c,** ChIP-seq experiments reveal recruitment of dCas9 to the target site (blue triangle) with a targeting ωRNA shown as two independent replicates. Data shown as in **a.** Analysis of off-target sites reveal a short (3–4 nt) seed sequence adjacent to the PAM motif. **d,** ChIP-seq experiments reveal recruitment of dCas12a to the target site (blue triangle) with a targeting ωRNA shown as two independent replicates. Data shown as in **a.** Analysis of off-target sites reveals a short (4–5 nt) seed sequence adjacent to PAM motif.

**Extended Data Figure 7.**
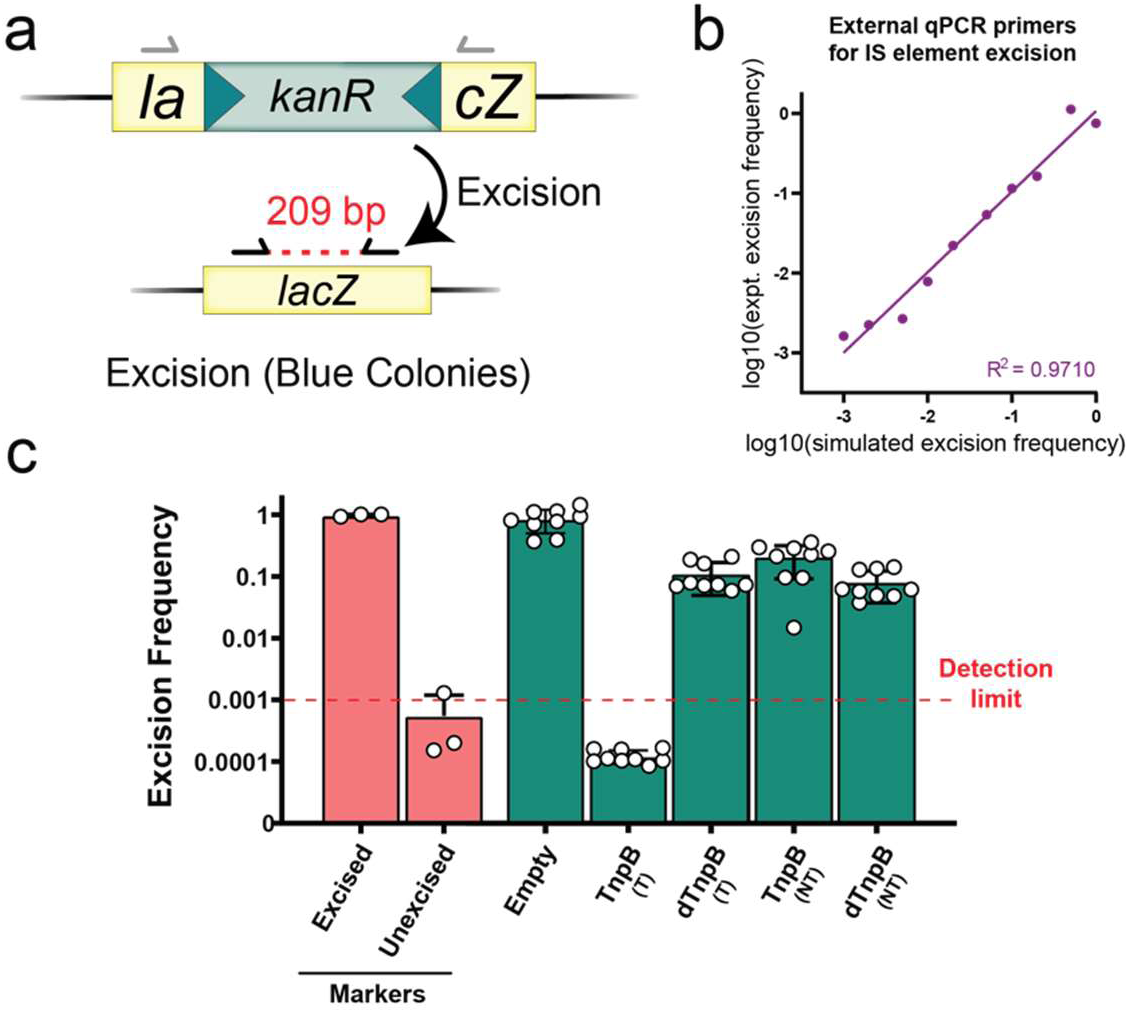
qPCR analysis of IS element loss upon TnpA and TnpB co-expression. **a,**Schematic of qPCR-based strategy for quantifying excision. Primers are designed to flank the donor joint following excision and religation. Selective PCR conditions with a shortened extension time allows for reduced amplification of the starting locus containing the mini-Tn. **b,** Comparison of simulated excision frequencies, generated by mixing clonally excised and unexcised lysate in known ratios, versus experimentally determined integration efficiencies measured by qPCR. **c,** qPCR-based quantification of transposon excision. Excised represents wild-type *lacZ* and unexcised represents lacZ containing mini-Tn as controls. TnpA was provided in all conditions shown in green. The detection limit is based on simulated excision frequencies shown in **b**.

