## Supplementary Figures for "Transposon-encoded nucleases use guide RNAs to selfishly bias their inheritance"

**Supplementary Figure 1 | Multiple sequence alignment of transposon left ends.**

Transposon boundary is delineated by dotted line. Transposon family is shown as indicated on right. Percent similarity is indicated at each position by color. Dark green represents 100% similarity. Light green represents 80-99% similarity and yellow represents 60-79% similarity.

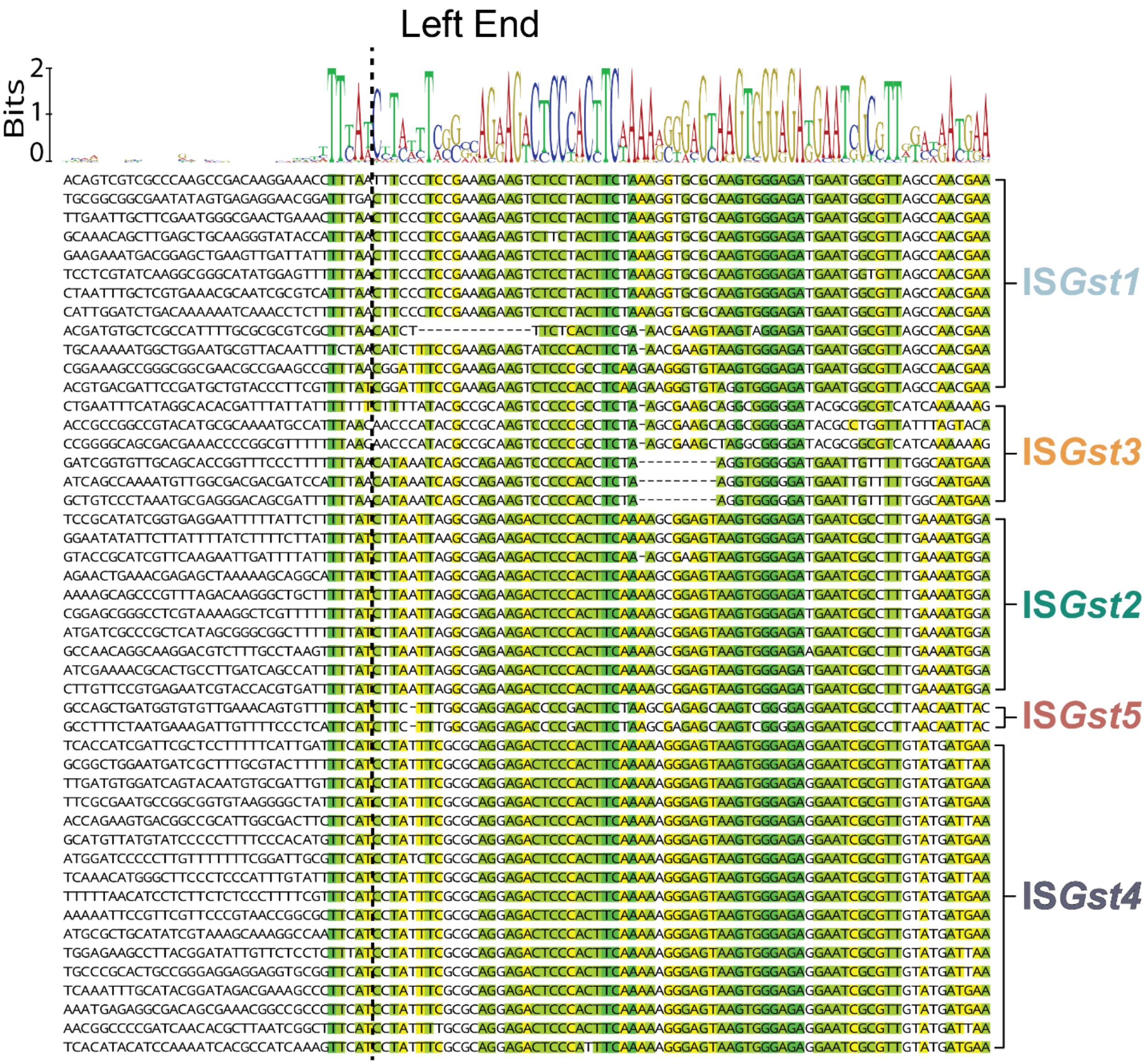

**Supplementary Figure 2 | Multiple sequence alignment of transposon right ends.**

Transposon boundary is delineated by dotted line. Transposon family is shown as indicated on right. Percent similarity is indicated at each position by color. Dark green represents 100% similarity. Light green represents 80-99% similarity and yellow represents 60-79% similarity.

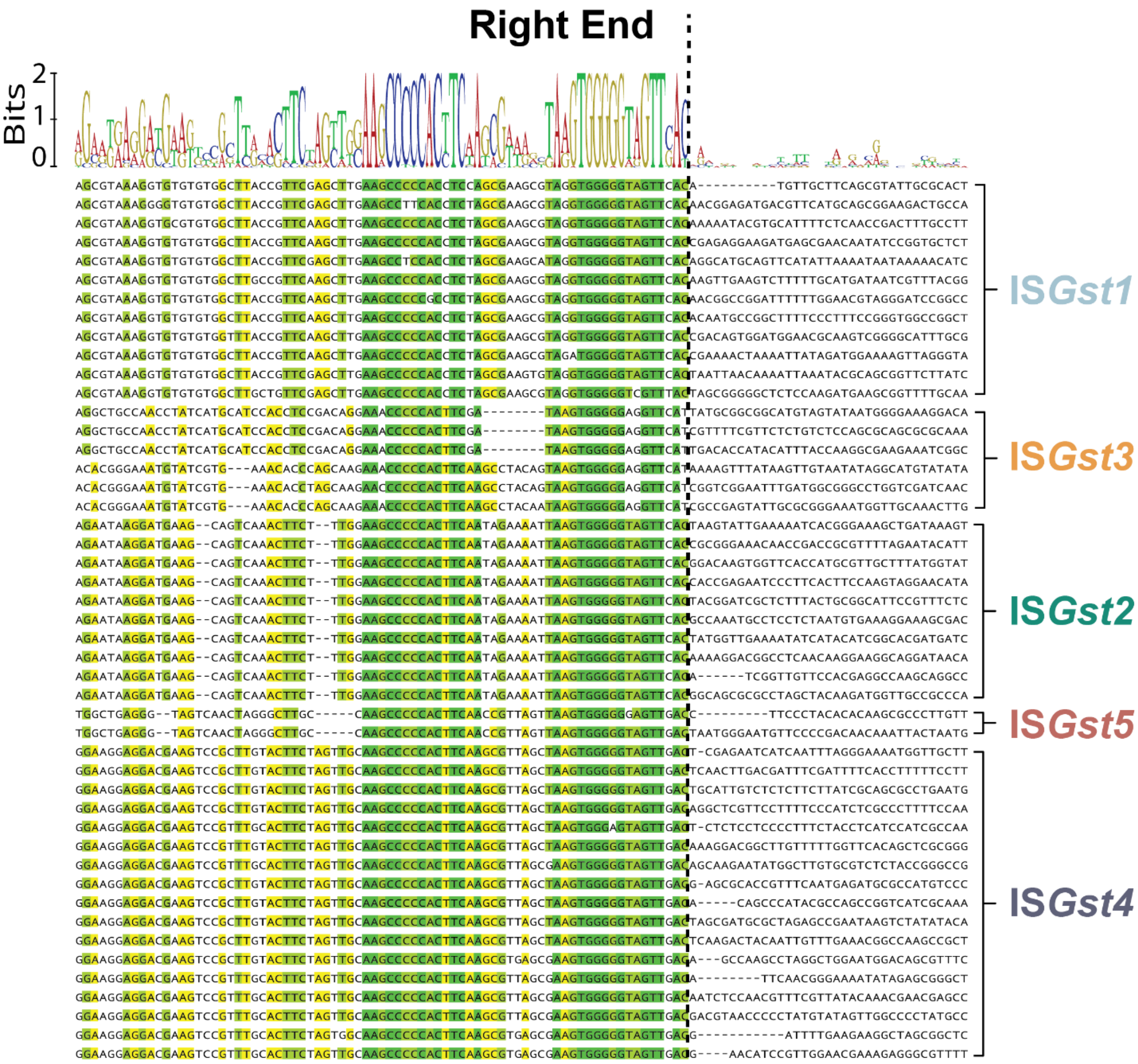
